# Machine learning-based definition of cellular senescence reveals pro-senescence potential implication in lung adenocarcinoma

**DOI:** 10.1101/2025.10.11.681821

**Authors:** Lifei Ma, Huiyang Li, Yiling Li, Zhi-Ai Lin, Jia-Qi Li, Yu-Zi Zhang, Peng Zhang, Zehao Yao, Jialing Li, Min Xiong, Yinghao Cao, Ruidong Li, Chen Yang, Xiaoqiang Tang, Minjiang Chen, He-Ping Wang, Wenjie Zheng, Jing Yang, Xiaoyue Wang, Gong-Hong Wei, Xiaoman Wang, Hou-Zao Chen

**Author notes:** Correspondence (Hou-Zao Chen); (Xiaoman Wang); (Gong-Hong Wei). These authors contributed equally: Lifei Ma and Huiyang Li.

## Abstract

Despite growing evidence implicating cellular senescence in tumor progression, methodological challenges in objectively quantifying senescent cell burden across cancer types continue to limit mechanistic studies and clinical translation. To address this, we developed the Predictive Cellular Senescence Model (PreCSenM), a machine learning-based tool that assigns a CS score by integrating senescence-associated features across 888 samples. Benchmarking analyses confirmed that PreCSenM markedly outperforms existing methodologies in reliably predicting senescent states within both normal and cancerous transcriptomic datasets. In lung adenocarcinoma (LUAD), PreCSenM identified the CS score as a robust predictor of clinical outcomes. Multi-omics analyses further revealed that lower CS levels correlate with decreased genomic stability and enhanced immune responses, aligning with clinical observations. Notably, drug discovery using PreCSenM identified histone deacetylase inhibitors (HDACis) as potent inducers of CS in LUAD. Transcriptional and epigenetic profiling of HDACi-treated cells pinpointed FOSB as a core transcription factor driving CS, and FOSB knockdown blocked HDACi-induced senescence. Overall, our study establishes PreCSenM as a multidimensional senescence quantification tool, bridging computational prediction with clinical relevance and mechanistic validation to enable senescence-directed therapies in precision oncology.

## Introduction

Cellular senescence (CS) refers to a stable state of cell cycle arrest accompanied by profound alterations in gene expression^1–4^. Recent studies have revealed that CS in cancer is highly context-dependent, varying across cancer types and tumor stages^4^. On the one hand, it serves as a natural barrier to tumorigenesis by inhibiting cell division and growth through epigenetic perturbations and immune-mediated clearance^3–6^. On the other hand, senescent cells secrete cytokines in a paracrine or endocrine manner, potentially creating a chronic inflammatory microenvironment that promotes tumorigenesis^7,8^. These findings highlight the importance of systematically evaluating CS burden in different cancer contexts, which could pave the way for tailoring senescence-based therapies across diverse cancer types.

Robust quantification of CS levels in cancer remains lacking. CS was originally defined based on canonical biological biomarkers such as senescence-associated β-galactosidase (SA-β-gal) activity and the cell cycle regulators p21 (CDKN1A) and p16 (CDKN2A) in non-malignant systems^1,9,10^. While these markers have advanced our understanding of CS, reliance on individual markers is inadequate for definitively assessing senescence levels^11^, particularly in oncological contexts. Tumor biology introduces unique complexities, as recurrent mutations in senescence-associated genes (e.g., TP53, CDKN2A) can compromise marker specificity^12^. Given the high inter- and intra-tumor heterogeneity in cancer, canonical senescence markers derived from non-malignant systems often exhibit limited specificity and context-dependent reliability in tumor settings^13^. Their expression does not always faithfully reflect true senescent states in cancer^11^. Therefore, the development of a robust and comprehensive senescence signature capable of capturing conserved features across both normal and cancerous cells is essential. An ideal biomarker for cancer-associated senescence should exhibit consistent expression both within individual tumors and across different tumor types, ensuring its reliability as a diagnostic and prognostic tool. To address these challenges, a promising strategy involves leveraging multigene panels derived from senescent normal cells and applying them to cancerous tissues. This approach could overcome the limitations of context-specific marker variability, offering a more accurate and transferable method for identifying senescence-associated features in the tumor microenvironment.

CS quantification has seen significant methodological advances, with established approaches including algorithmic metrics such as hUSI and SenCID^14–17^, as well as transcriptional signatures^18–24^, which outperformed traditional senescence-associated markers, such as TP53 and CDKN2A, in distinguishing senescent cells. However, most of these tools lack a universal and streamlined framework capable of translating CS levels into clinical applications across cancers or generalizing to broader biological contexts. Many existing models require disease-specific parameter tuning and technical expertise, limiting their accessibility and widespread use. Given the strong predictive power of machine learning algorithms in biomarker discovery^25,26^, integrating diverse machine learning strategies into CS quantification offers the potential to uncover novel senescence-associated biomarkers and therapeutic targets in a context-independent and user-friendly manner^27,28^.

In this study, we present the Predictive Cellular Senescence Model (PreCSenM), a novel computational approach designed to quantify CS levels. By leveraging multiple machine learning algorithms, we analyzed multi-platform transcriptomic data from 888 cell lines and tissue samples, encompassing over 13 distinct normal senescent tissue types and more than 10 cancer types. This comprehensive analysis enabled the construction of a consensus cellular senescence-related gene signature (CSGS). Using this framework, we identified associations between senescence levels, molecular characteristics, and clinical outcomes. We further examined the effects of small-molecule histone deacetylase inhibitors (HDACis) on senescence levels in lung adenocarcinoma (LUAD) cells. Transcriptomic and epigenomic profiling of HDACi-treated cells revealed that FOSB modulates the senescence program. Our integrated analyses establish a machine learning–based framework for effectively evaluating senescence levels in cancer and provide insights into the biological regulation and translational potential of CS in LUAD. To support further research and clinical translation, we developed a user-accessible web portal (http://precsenm.bmicc.org/) that enables senescence prediction across diverse contexts.

## Results

### Consensus gene signatures for CS activity

Given the lack of universal and specific markers to identify and examine CS, we applied machine-learning approaches to classify senescent cells based on gene expression profiles (Fig. 1). We collected multi-platform expression profiles from 519 samples across more than 13 types of senescent tissues, including both senescent and non-senescent normal cell lines, and 369 samples from over 10 cancer types in tumor cell lines or tissues, totaling 888 samples. These samples include various senescence-inducing conditions such as replicative, DNA damage, gene editing, oncogene, oxidation, and others. (Extended Data Fig. 1a, b, Supplementary Table 1, 2). Approximately 70% of all normal senescent cell lines were randomly assigned to the training set, with the remaining 30% used for testing. After rigorous preprocessing to remove potential confounding factors, we used learning-based methods to extract 236 genes for constructing CSGS based on the training data. The signature showed enrichment in senescence-related biological processes, including cell cycle, mRNA processing, RNA splicing, cellular response to DNA damage stimulus, and mRNA metabolic process (Fig. 2a). These genes include several well-established senescence-associated genes (*CDKN1A*, *CCND1*, *LMNB1*, *HMGB1*, and *MCM7*), those involved in mRNA processing and splicing (*SNRPB2*, *SRSF1*, and *SRSF7*), interferon response genes (*IFI6*), and histone variants (*H2AC6*, *H2AZ2*, and *H2BC21*). Taken together, the CSGS based on multi-platform data likely reflects well-established characteristics of CS.

**Fig. 1.**
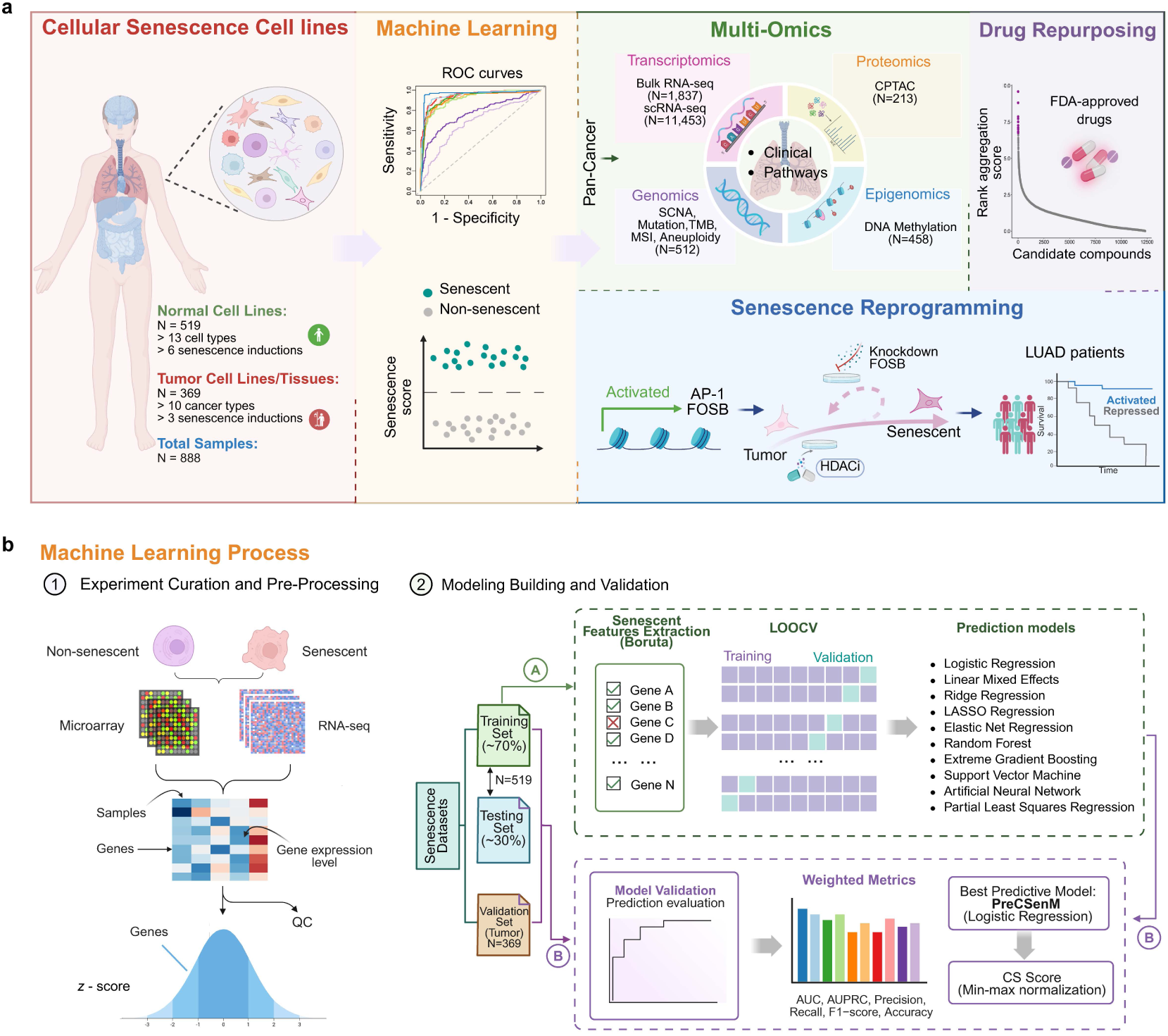
Overview of the development and application of the PreCSenM. **a** Schematic diagram illustrating the development of the PreCSenM computational model for evaluating cellular senescence (CS) levels, along with its application in lung adenocarcinoma (LUAD). Applications include multi-omics analyses, drug repurposing strategies, and investigation of senescence programming mechanisms. **b** The workflow consists of two main steps: (1) Experimental curation and pre-processing, including the identification and collection of 888 publicly available RNA-seq and microarray datasets from normal and tumor cell lines or tissue samples, quality control assessment, and computation of z-scores for each dataset; (2) Model building and validation, including senescence feature extraction using the Boruta algorithm to identify a key cellular senescence-related gene signature (CSGS), distinguishing senescent from non-senescent states, constructing ten machine learning models based on the CSGS signature, and evaluating model performance using multiple metrics. Further methodological details are provided in the Methods section.

**Fig. 2.**
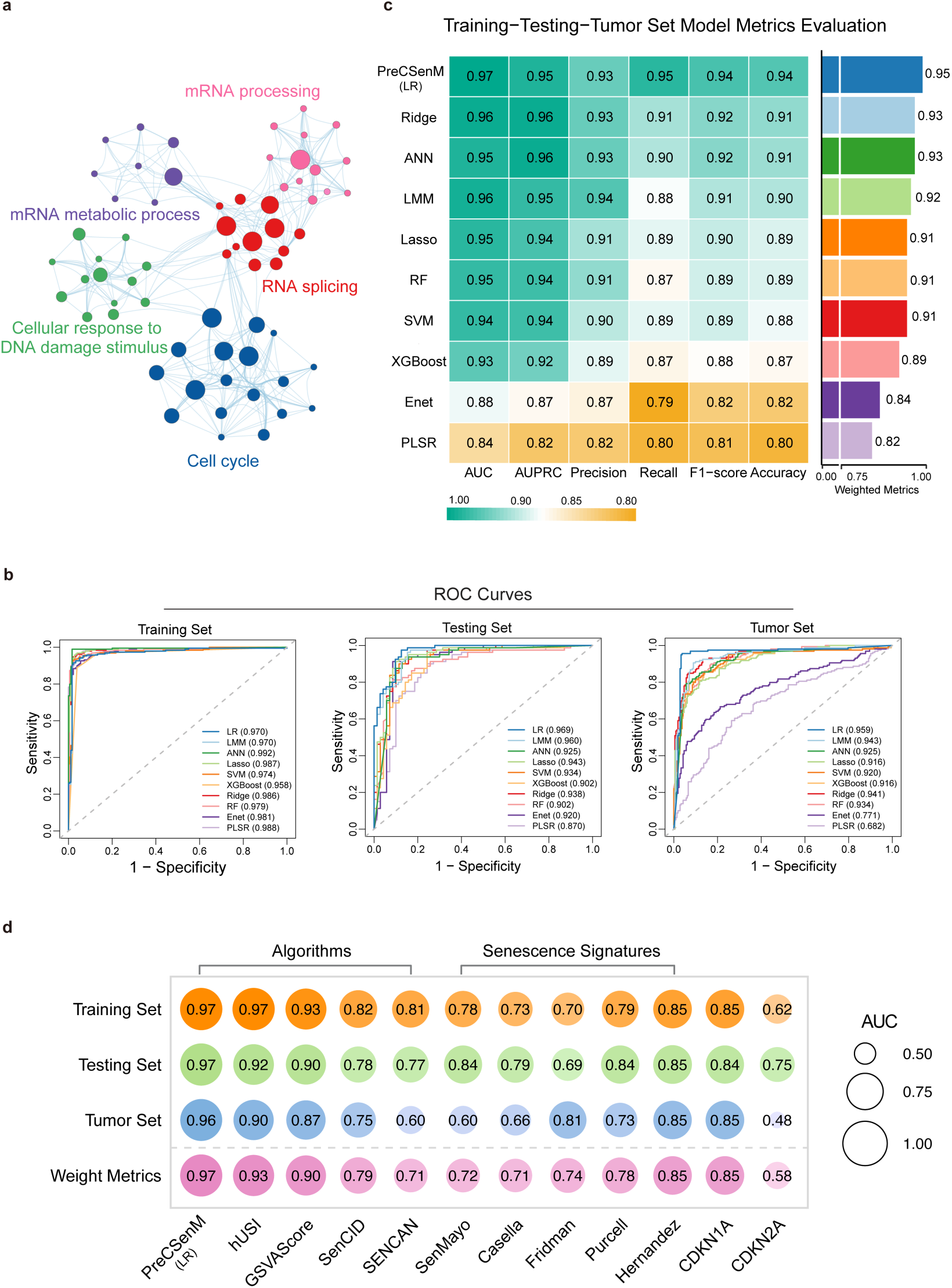
Performance evaluation of PreCSenM across diverse datasets and algorithms. **a** Five color-coded core ontology clusters represent the core enrichment network of CSGS. The network was visualized using Cytoscape software. **b** Receiver operating characteristic (ROC) curves and corresponding area under the curve (AUC) values for ten machine learning models were evaluated across three datasets: the training set (left) and testing set (middle), comprising a total of 519 samples spanning 13 diverse normal senescent tissue types, and the tumor set (right), which includes samples from over 10 different cancer types. AUC values were averaged over 10 repeated runs, each assessing model sensitivity across varying false positive rates (1 − specificity) through ROC curve analysis. **c** Performance of ten predictive model types was evaluated using accuracy, precision, recall, F1 score, area under the receiver operating characteristic curve (AUC), and area under the precision-recall curve (AUPRC) across training, testing, and validation (tumor) datasets. A barplot visualization compares the weighted scores of these seven metrics to assess the overall performance of the ten machine learning models. Weighted scores were calculated by integrating all metrics, adjusted for sample size. **d** Benchmarking of PreCSenM was performed against individual canonical markers (CDKN1A and CDKN2A), previously published senescence signatures, and existing algorithms for senescence classification across three datasets, with weighted values used for comparison. **Abbreviations:** LR, logistic regression; LMM, linear mixed-effects model; ANN, artificial neural network; Lasso, LASSO regression; SVM, support vector machine; XGBoost, extreme gradient boosting; Ridge, ridge regression; RF, random forest; Enet, elastic net regression; PLSR, partial least squares regression.

### PreCSenM as a robust classifier for senescent cells in tumor contexts

To develop an effective model for predicting cellular senescence, we evaluated ten classification methods using the CSGS, including linear mixed-effects models (LMM), logistic regression (LR), ridge regression (Ridge), Lasso regression (Lasso), elastic net regression (Enet), random forest (RF), extreme gradient boosting (XGBoost), support vector machine (SVM), artificial neural networks (ANN), and partial least squares regression (PLSR) through leave one out cross validation (LOOCV) in the training dataset to determine optimal parameters (see Methods). Performance metrics were comparable across training, testing, and validation (tumor) datasets for all models, except for Enet and PLSR (Fig. 2b, c; Extended Data Fig. 1c). This suggests that the models effectively generalized to unseen data, demonstrating robust performance on both testing and tumor datasets. The optimal LR model included all metrics with the highest weighted index (0.95) in the training, testing and tumor datasets (Fig. 2c). Furthermore, the performance of PreCSenM was not significantly confounded by data types, senescence-induced types or cell types (Extended Data Fig. 1d). Additionally, we applied PreCSenM model to previously published scRNA-seq data with senescent status^29^. Senescent status cells consistently displayed significantly higher CS scores than non-senescent status cells (Extended Data Fig. 1e). Building on these findings, we developed PreCSenM, which leverages the CSGS and optimal LR model coefficients to robustly and accurately distinguish senescent from non-senescent cells in both normal and tumor contexts.

To assess the performance of the PreCSenM classifier in predicting senescent cells, we compared it against single canonical markers (CDKN1A and CDKN2A)^30,31^ and five publicly available senescence signatures (SenMayo, Casella, Fridman, Purcell and Hernandez)^18–24^ as well as four senescence gene scoring algorithms (GSVAscore, SenCID, hUSI and SENCAN)^14–17^ (Supplementary Table 3). The area under the curve (AUC) values were evaluated among the single markers, senescence signatures, gene scoring algorithms, and the PreCSenM method across training, testing, and tumor datasets (Fig. 2d). PreCSenM demonstrated superior performance across all cohorts, achieving the highest weighted AUC of 0.97, outperforming all other methods and algorithms (Fig. 2d).

### CS score from PreCSenM as a potential predictor of clinical outcomes in LUAD

Existing evidence revealed that in most cancer types, the senescence level is lower in tumor tissues than in normal tissues^18^. We then applied the CS scores to a cohort from The Cancer Genome Atlas (TCGA) database comprising ten cancer types with more than 35 samples. Consistently, normal tissue exhibited a higher senescence level than tumor tissue (Fig. 3a), except for thyroid cancer (THCA) tissue, as previously reported^18^. These results further confirmed that the PreCSenM classifier can detect the senescent status in different tumor tissues.

**Fig. 3.**
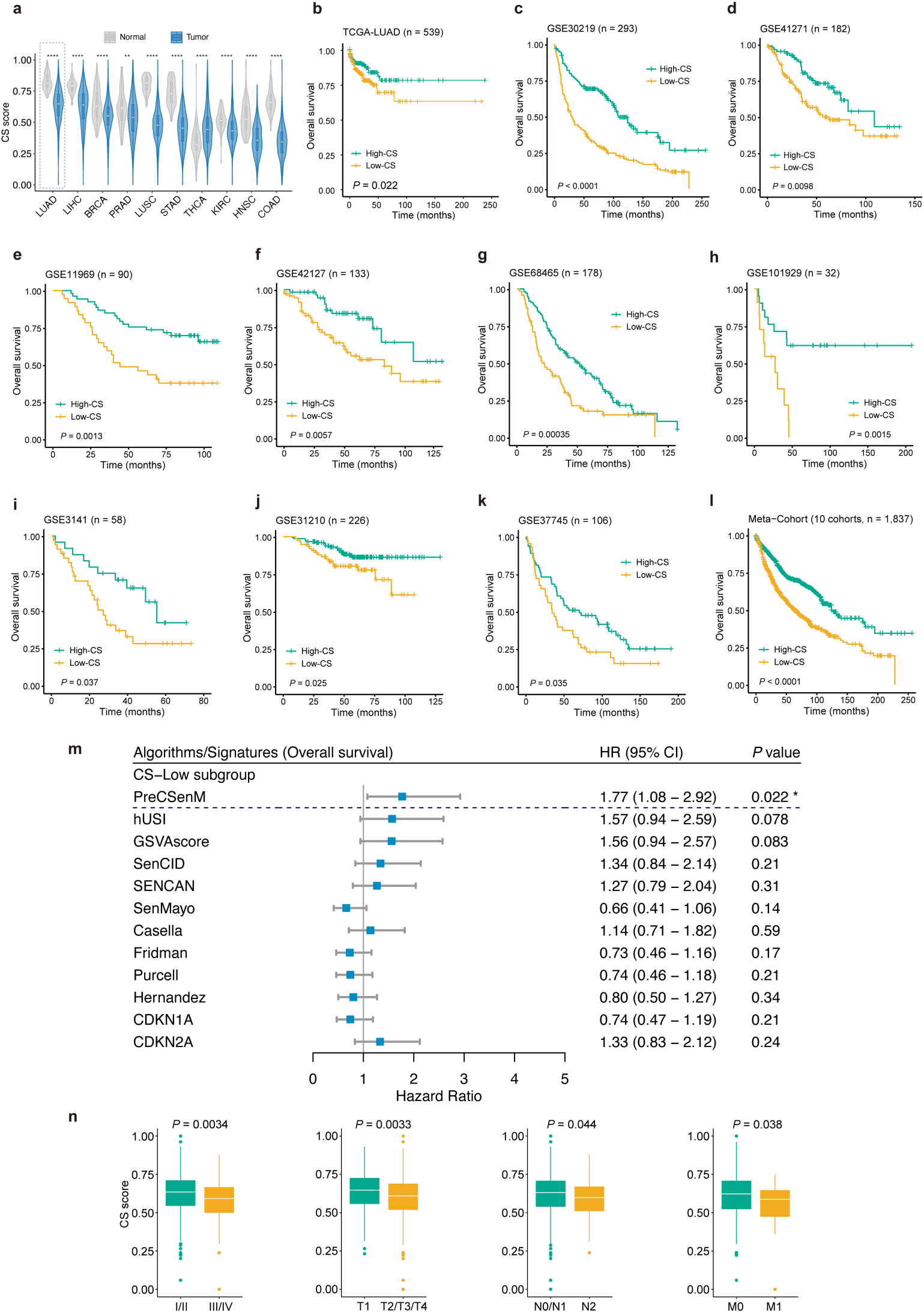
PreCSenM-derived cellular senescence (CS) score showed significant correlations with clinical indices in patients with lung adenocarcinoma (LUAD). **a** CS scores in primary tumors (blue) compared with those in adjacent normal solid tissues (gray). A total of 10 cancer types from The Cancer Genome Atlas (TCGA) dataset with > 35 normal and tumor samples were selected, which were ordered by decreasing the distribution of the CS score of tumors. \*\*\*\**P* < 0.0001, \*\**P* < 0.01 (Wilcoxon test). **b–l** Kaplan–Meier curves of overall survival (OS) according to the CS score subgroups in TCGA-LUAD (log-rank test: *P* = 0.022, *n* = 539) (b), GSE30219 (log-rank test: *P* < 0.0001, *n* = 293) (c), GSE41271 (log-rank test: *P* = 0.0098, *n* = 182) (d), GSE11969 (log-rank test: *P* = 0.0013, *n* = 90) (e), GSE42127 (log-rank test: *P* = 0.0057, *n* = 133) (f), GSE68465 (log-rank test: *P* = 0.00035, *n* = 178) (g), GSE101929 (log-rank test: *P* = 0.0015, *n* = 32) (h), GSE3141 (log-rank test: *P* = 0.037, *n* = 58) (i), GSE31210 (log-rank test: *P* = 0.025, *n* = 226) (j), GSE37745 (log-rank test: *P* = 0.035, *n* = 106) (k), and meta-cohort (log-rank test: *P* < 0.0001, *n* = 1,837) (l). **m** Cox proportional hazards model and survival analysis of CS evaluation methods in TCGA LUAD. The forest plot displays hazard ratios and 95% confidence intervals (CIs) for overall survival in CS subgroups, with significance defined as log-rank *P* < 0.05. **n** CS scores across American Joint Committee on Cancer (AJCC) and TNM stages in TCGA-LUAD patients. AJCC stages: I/II vs. III/IV (*n* = 421 vs. 110); T stages: T1 vs. T2/T3/T4 (*n* = 176 vs. 360); N stages: N0/N1 vs. N2 (*n* = 447 vs. 74); M stages: M0 vs. M1 (*n* = 365 vs. 25). Wilcoxon test P-values were reported for comparisons between groups.

Lung cancer has one of the lowest 5-year survival rates among cancer types and stands as the primary cause of cancer-related deaths^32^. Numerous accessible multi-omics data resources for LUAD are publicly available. The CS level of LUAD was the highest among the ten types of cancer (Fig. 3a). To investigate the relationship between the CS score and clinicopathological characteristics, we collected ten LUAD datasets, and a total of 1,837 patients were stratified into high- and low-CS groups based on the optimal cut-off value determined independently for each dataset (Extended Data Fig. 2a). Survival analysis revealed that patients in the high-CS group had significantly better overall survival (OS) compared to those in the low-CS group across all datasets (Fig. 3b–k). The meta-cohort comprising all 1,837 samples showed the same tendency (*P* < 0.0001; Fig. 3l). Furthermore, we performed a comparative survival analysis using various CS evaluation methods within the TCGA LUAD dataset. PreCSenM consistently outperformed other methods, yielding statistically significant correlations with OS, thus highlighting its robustness and potential clinical utility (Fig. 3m). Additionally, high CS levels were significantly correlated with other clinicopathological characteristics, including vital status, tumor pathological American Joint Committee on Cancer (AJCC) stages, and T, N, and M stages (Fig. 3n, Extended Data Fig. 3a–i). Tumors exhibiting high levels of CS generally have lower tumor purity (Extended Data Fig. 2b, c), which is consistent with the increased infiltration of stromal or immune cells within senescence-enriched tumor microenvironments^15^. Additionally, tumor purity showed no significant impact on survival outcomes in LUAD (Extended Data Fig. 2d). These results suggest that the CS score may serve as an independent marker for predicting prognosis in LUAD.

### Relationships between multi-omic molecular characteristics and CS in LUAD

To gain insights into the clinical significance of CS in LUAD, we applied multi-omic data to evaluate correlations between CS score and omic features in TCGA-LUAD data. At the genomic level, we examined mutation profiles of key senescence-related genes in LUAD cohorts stratified by high- and low-CS groups. The analysis revealed a significantly elevated mutation burden in most genes among low-CS patients compared to their high-CS counterparts (Fig. 4a, b). Notably, *TP53*—a pivotal regulator of the cell cycle/TP53 pathway—showed the most pronounced disparity in mutation frequency in low-CS patients (59% in low-CS vs. 37% in high-CS; *P* = 6.27e-08), while *RB1* mutations were also enriched in low-CS patients (8% vs. 2%; *P* = 3.75e-04). These results underscore a pronounced prevalence of *TP53* mutations in low-CS patients, consistent with their significant association with clinically aggressive tumor behavior^3–6^. In addition, we measured mutations by somatic copy-number alteration (SCNA) score^33^, the sum of alterations at the focal, arm, and chromosome levels. In LUAD patients, the overall SCNA score significantly decreased with the CS score (R^2^ = −0.35, *P* = 2.3e-16; Fig. 4c). Consistently, we observed a stronger negative association between CS score and tumor mutation burden (TMB), aneuploidy score (AS) and microsatellite instability (MSI) score (Extended Data Fig. 4a–c). These results demonstrate that genomic alterations are significantly decreased in high-CS patients with LUAD, consistent with the better clinical outcome of patients with high-CS levels (Fig. 2).

**Fig. 4.**
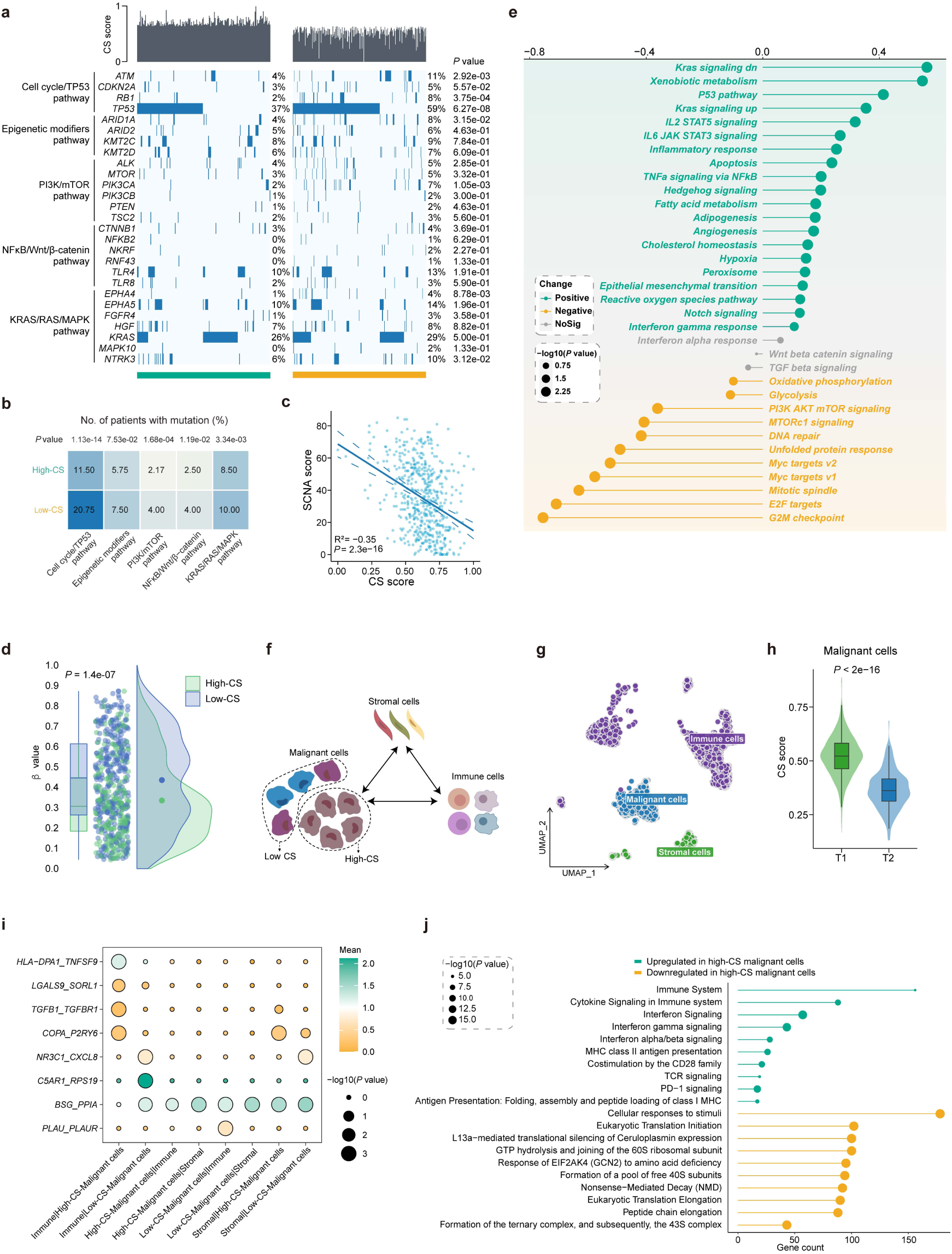
PreCSenM-derived CS score identified multi-omic molecular characteristics in patients and malignant cells with lung adenocarcinoma (LUAD). **a** Oncoprint of the mutation status of genes in various senescence-related pathways involved in LUAD. Each column represents a sample, blue boxes indicate non-synonymous mutations, and each bar (upper panel) shows the CS score of the patients. Fisher’s exact test was used to establish statistical significance. **b** Number of mutations in different pathways and distinct subgroups. *P* values were calculated using Fisher’s exact test. **c** Correlations between somatic copy-number alteration (SCNA) and CS scores in LUAD (rank-based Spearman’s correlation). **d** Box and ridge plots showing the distribution of estimated β values between the two senescence subgroups based on the LUAD database. Dots in the ridge plot represent the median values for each group. The Wilcoxon test was used to calculate *P* values. **e** Spearman correlation between CS scores and pathway activities. Pathway activity was estimated using gene set variation analysis (GSVA) and represented by normalized enrichment scores (NES) for individual TCGA-LUAD patients. The dot size denotes the statistical significance of the correlation, and the colors indicate the direction of association—green for positive correlations, yellow for negative correlations, and gray for non-significant results. **f** The design of the LUAD scRNA-seq dataset enables the investigation of CS heterogeneity among malignant cells (high-CS or low-CS), and their relationships with two other major cell types within the tumor microenvironment. **g** Uniform Manifold Approximation and Projection (UMAP) plot displaying cells colored by the three major cell types at the single-cell level in LUAD. Each point represents a single cell. **h** Violin plots showing CS scores across different T stages in malignant cells from LUAD scRNA-seq data. Wilcoxon test *P* values are presented. **i** Bubble graphs illustrating significant ligand–receptor interactions between high-CS/low-CS malignant cells and neighboring cells. Ligand- and receptor-expressing cells are shown on the x-axis, and ligands and receptors on the y-axis. Bubble size denotes the importance of interactions between the ligand and receptor, and color represents the average expression levels in the interacting cells (permutation test in CellPhoneDB). Significant predicted interaction pairs are those with an average expression > 0.2 and *P* < 0.05. **j** Reactome pathway analysis of differentially expressed genes between the two CS groups in malignant cells with the selected significantly enriched terms. The size of each point denotes the significance of enrichment. Upregulated pathways in the high-CS malignant cells are denoted in green, whereas downregulated pathways are denoted in yellow.

From the epigenetic aspect, we investigated senescence-related epigenetic changes in TCGA-LUAD patients. The level of DNA methylation of differentially methylated probes (DMPs) in two senescence subgroups was lower in the group with higher CS scores (Fig. 4d). Furthermore, the DMPs with strong signals exhibited negative relationships with senescence levels (e.g., *CPXM1*, a member of the CSGS; Extended Data Fig. 4d). These results show that patients with low senescence levels exhibit hypermethylation^34^.

Using transcriptomic data, we calculated the pathway activity of hallmark pathways by gene set variation analysis (GSVA) and found that 34 of 50 pathways were connected to senescence-associated transcriptomics in LUAD. Particularly, significant negative associations with the CS score were observed in proliferation-related gene sets (including MYC targets, G2M checkpoint, and E2F targets), as well as oxidative phosphorylation, MTORC1 signaling, DNA repair, and the unfolded protein response (Fig. 4e). Inflammatory responses (IL2 STAT5 signaling, IL6 JAK STAT3 signaling, TNFα signaling via NFκB, and interferon gamma response), as well as Kras signaling, the P53 pathway, and apoptosis, were significantly positively correlated with the CS score (Fig. 4e). Overall, transcriptomic analysis reveals that pathway alterations are associated with CS scores, indicating LUAD–specific senescence programs.

To further investigate the role of CS in lung cancer at the single-cell level, we carried out single-cell analyses following the standard MAESTRO workflow^35^. Cell clusters were mapped into three major cell types, including immune, stromal, and malignant cells. The malignant cell population was further divided into two subgroups: CS-high and CS-low (Fig. 4f, g). Additionally, the CS level of malignant cells was significantly higher in the T1 stage than T2 stage (Fig. 4h), which is consistent with previous results (Fig 3m). Furthermore, we investigated the altered cell–cell interactions with senescent levels. Certain ligand-receptor pairs (Fig. 4i; i.e., HLA−DPA1-TNFSF9, LGALS9−SORL1, TGFB1−TGFBR1, and COPA−P2RY6) were observed between immune cells and high-CS-malignant cells had a proinflammatory effect on cancer, while ligand–receptor pairs (i.e., C5AR1–RPS19, NR3C1–CXCL8, and BSG–PPIA) between immune cells and low-CS-malignant cells promoted cancer progression and metastasis. Pathway enrichment of differentially expressed genes (DEGs) of high- and low-CS malignant cells revealed the enrichment of immune-related pathways, including cytokine signaling in immune system, adaptive immune system, interferon signaling, and PD-1 signaling (Fig. 4j).

Collectively, these findings demonstrate the multi-omic features of PreCSenM-derived CS score in the LUAD patients, which may provide mechanistic explanation for the significant associations between CS score with clinical outcomes in LUAD.

### Identification of senescence-associated drugs using CS score

To advance the development of senescence-related drugs in lung cancer, we systematically identified potential drugs according to the following pipeline (Fig. 5a). First, we generated a CS- and survival-associated signature using a step-wise process in the TCGA-LUAD cohort. A total of 6,717 DEGs (adjusted *P* < 0.05 and |fold change FC| > 2) between the high- and low-CS groups were retained as senescent features of LUAD for further analysis. Second, we conducted univariate Cox regression analyses to determine the prognostic relevance of these DEGs for OS. We found that 205 were significantly correlated with OS. Third, we narrowed down the size of the query signature to the recommended size. To achieve this, we constructed a random survival forest (RSF) model with 1000 trees grown each time and repeated 1000 times. Finally, 182 genes with positive variable importance values were included (Fig. 5a).

**Fig. 5.**
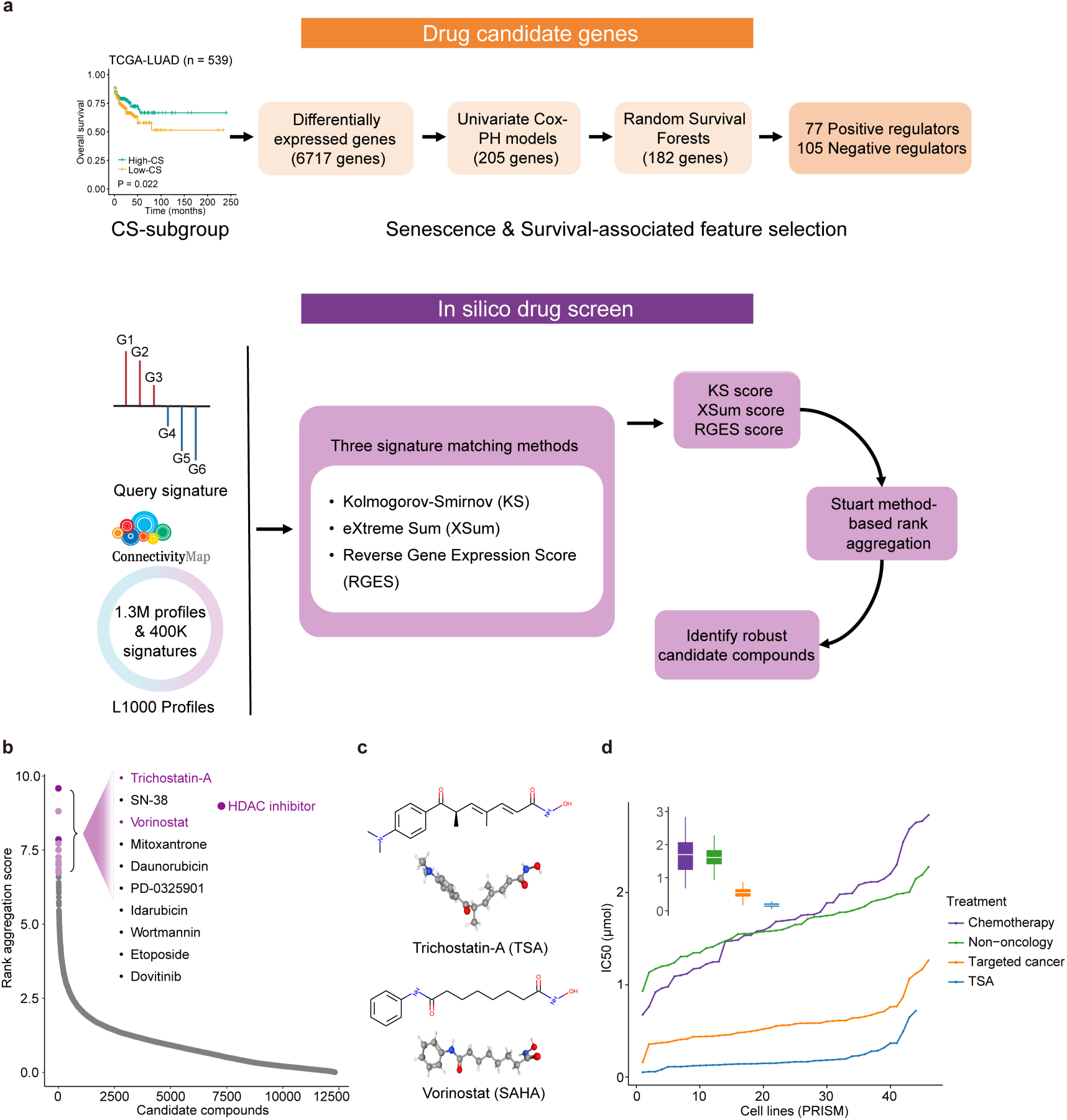
Overview of data-driven therapeutic discovery by PreCSenM. **a** Generation of optimal query signatures associated with CS and patient survival for drug screening via a step-wise process in the CGA-LUAD dataset. This process involved differential expression analysis between the high- and low-CS groups, univariate Cox proportional hazard regression analyses, and random survival forest selection (top panel). CS-related candidate drugs were identified by querying CS- and survival-associated gene signatures against the LINCS database, followed by prioritization using three distinct signature-matching approaches to select robust candidates (bottom panel). **b** Ranking results of drug predictions, with the top ten drugs labeled. **c** Two- and three-dimensional chemical structures of trichostatin-a (TSA; top) and vorinostat (SAHA; bottom). **d** TSA and three other pharmacological categories–chemotherapy, targeted cancer treatment, and non-oncology–were compared in terms of the distribution of compound activity. The median IC50 values of all substances in each category of the LUAD cell line were calculated using the PRISM dataset.

Based on these 182 genes, we performed signature matching analysis with three complementary methods, including Kolmogorov–Smirnov (KS)^36^, eXtreme Sum (XSum)^37^, and the Reverse Gene Expression Score (RGES)^38^ in the context of the A549 lung cancer cell lines from the Library of Integrated Network-Based Cellular Signatures (LINCS) program (Fig. 5a). The results were integrated using the order statistics-based method (Fig. 5b and Supplementary Table 4) for a rank aggregation analysis to produce a robust drug prediction^39^. Notably, HDACi, including trichostatin-a (TSA) and vorinostat (SAHA), exhibited the most significant potency among all drug candidates (Fig. 5b, c). Due to the extensive research of TSA data in the LUAD cell line, we focused on comparing IC_50_ values of TSA with other therapeutic pharmacological categories, including chemotherapy, targeted cancer agents, and non-oncology. The IC_50_ values of TSA were lower than other pharmacological categories (Fig. 5d), suggesting a higher drug sensitivity of the LUAD cell line to TSA treatment.

### Predicted drug (HDACi) induce LUAD cell senescence

To test whether HDACi can induce senescence in LUAD cells, we performed RNA-seq of A549 treated with TSA at different time points (Fig. 6a). Notably, treatment with HDACis reduced the viability of cancer cells over time (Fig. 6b). We assessed the senescence levels using CS scores from PreCSenM and found constantly increased CS scores over treatment time, particularly from 6–12 h (Fig. 6c). Differential gene expression analysis showed that the largest number of DEGs appeared at 12 h (Fig. 6d). Moreover, Gene Ontology (GO) analyses of DEGs revealed significant enrichment in senescence-related biological functions, such as cell cycle arrest, cell cycle G2/M phase transition, and histone modification at 12 h (Fig. 6e).

**Fig. 6.**
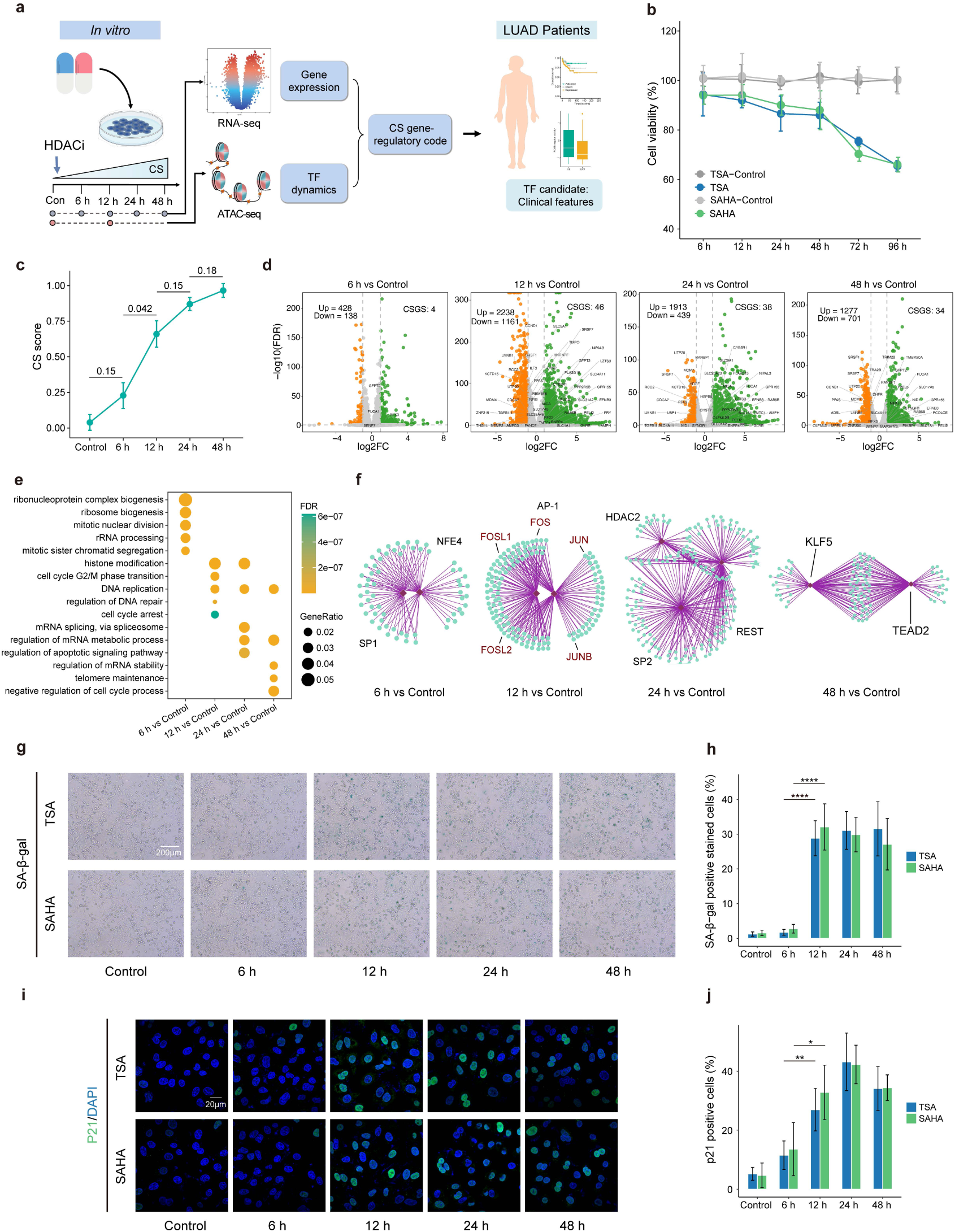
Validation of histone deacetylase inhibitor (HDACi) as a senescence-inducing drug and establishment of a senescence transcriptional program. **a** Schematic overview of defining the senescent features and identifying the core transcription factors (TFs) by analysis of RNA-seq data and assay for transposase-accessible chromatin with sequencing (ATAC-seq) data of A549 cells treated with histone deacetylase (HDAC) inhibitor at different time points (left). A relationship between core TFs and clinical features was identified in patients from The Cancer Genome Atlas (TCGA)-lung adenocarcinoma (LUAD) dataset (right). **b** Viability curves depicting the percentage of viable A549 cells incubated with TSA or vorinostat (SAHA) at their optimal concentrations (300 and 500 nM, respectively) for 96 h, with the control as the reference. **c** Senescence levels evaluated with PreCSenM using the RNA-seq data from A549 cells treated with trichostatin-a (TSA). Statistical differences between groups were determined using Student’s *t*-test. Data are presented as the mean ± SD. **d** Volcano plots showing differentially expressed genes (DEGs) at 6, 12, 24, and 48 h compared with the control. Green dots: upregulated genes, yellow dots: downregulated genes (FDR < 0.05, log2[fold change] ≥ 1 or ≤ –1). Cellular senescence-related gene signatures (CSGS) are labeled, and the number of DEGs and CSGSs was quantified. **e** Selected biological process Gene Ontology (GO) terms are enriched in each comparison, with the top five significantly enriched terms shown. The dot size corresponds to the number of genes in the GO term, and the color denotes enrichment significance. **f** TF enrichment was analyzed using RcisTarget based on DEGs from different comparisons. Only the most significantly enriched TFs are shown. AP-1 family TFs are highlighted during drug treatment for 12 h. **g, h** Representative senescence-associated β-galactosidase (SA-β-gal) staining images (g) of A549 cells treated with TSA (300 nM) or SAHA (500 nM) at different time points and quantitation of SA-β-gal+ senescent cells (h). *n* = 6 biologically independent samples, with three randomly selected regions per sample analyzed. Scale bar, 200 μm (Student’s *t*-test), and only the common groups with significant differences are shown. Data are represented as the mean ± SD. \*\*\*\**P* < 0.0001. **i, j** Representative images (i) of p21 immunofluorescence staining in A549 cells (p21, green; DAPI, blue) treated with TSA (300 nM) or SAHA (500 nM) at different time points, and quantification of stained cells (j); *n* = 6 biologically independent samples, with three randomly selected regions per sample analyzed. Scale bar, 20 μm (Student’s *t*-test), and only the common groups with significant differences are shown. Data are represented as the mean ± SD. \*\**P* < 0.01, \**P* < 0.05.

To confirm whether senescence was induced in the HDACi-treated A549 cells, we detected SA-β-gal activity, a well-established marker for CS. Consistent with CS score (Fig. 6c), SA-β-gal-positive cells significantly increased from 6 to 12 h when stimulated with TSA (300 nM) or SAHA (500 nM; Fig. 6g). Time-course kinetics analysis with different time points revealed that the pro-senescent potency of TSA and SAHA plateaus after a 12 h incubation (Fig. 6h). Given that p21 regulates the cell cycle and plays a crucial role in triggering senescence of cancer cells, we next examined whether TSA (300 nM) or SAHA (500 nM) could activate p21 in the nuclei of A549 cells. Immunofluorescence analysis showed significantly upregulated p21 protein levels in A549 cells treated with TSA or SAHA at 12 h (Fig. 6i, j). We further confirmed our result at 12 h by staining telomere-associated foci (TAF), makers of persistent DNA damage and cellular senescence^40,41^ (Extended Data Fig. 5a, b). These findings suggest that CS was induced in A549 cells by the HDACi after a 12 h incubation.

### The AP-1 family is involved in enhancing HDACi-induced senescence

Epigenetic mechanisms are important in CS, initiation and progression^42^. To reveal the chromatin accessibility dynamics of HDACi-induced senescence, we performed a time-course Assay for Transposase-Accessible Chromatin with sequencing (ATAC-seq) on TSA-treated A549 cells (Fig 7a). The 12 h drug treatment group displayed a higher ATAC peak signal in the promoter regions than the control. Hence, the chromatin progressively increased availability by 12 h (Fig. 7a). As the epigenetic status of promoter regions affects how downstream genes are expressed^43^, we conducted an integrative analysis of DEGs using RNA-seq and differential accessibility region (DAR) analysis with ATAC-seq; CS-associated up-regulated genes had significantly higher promoter accessibility than CS-associated down-regulated genes (Fig. 7b).

**Fig. 7.**
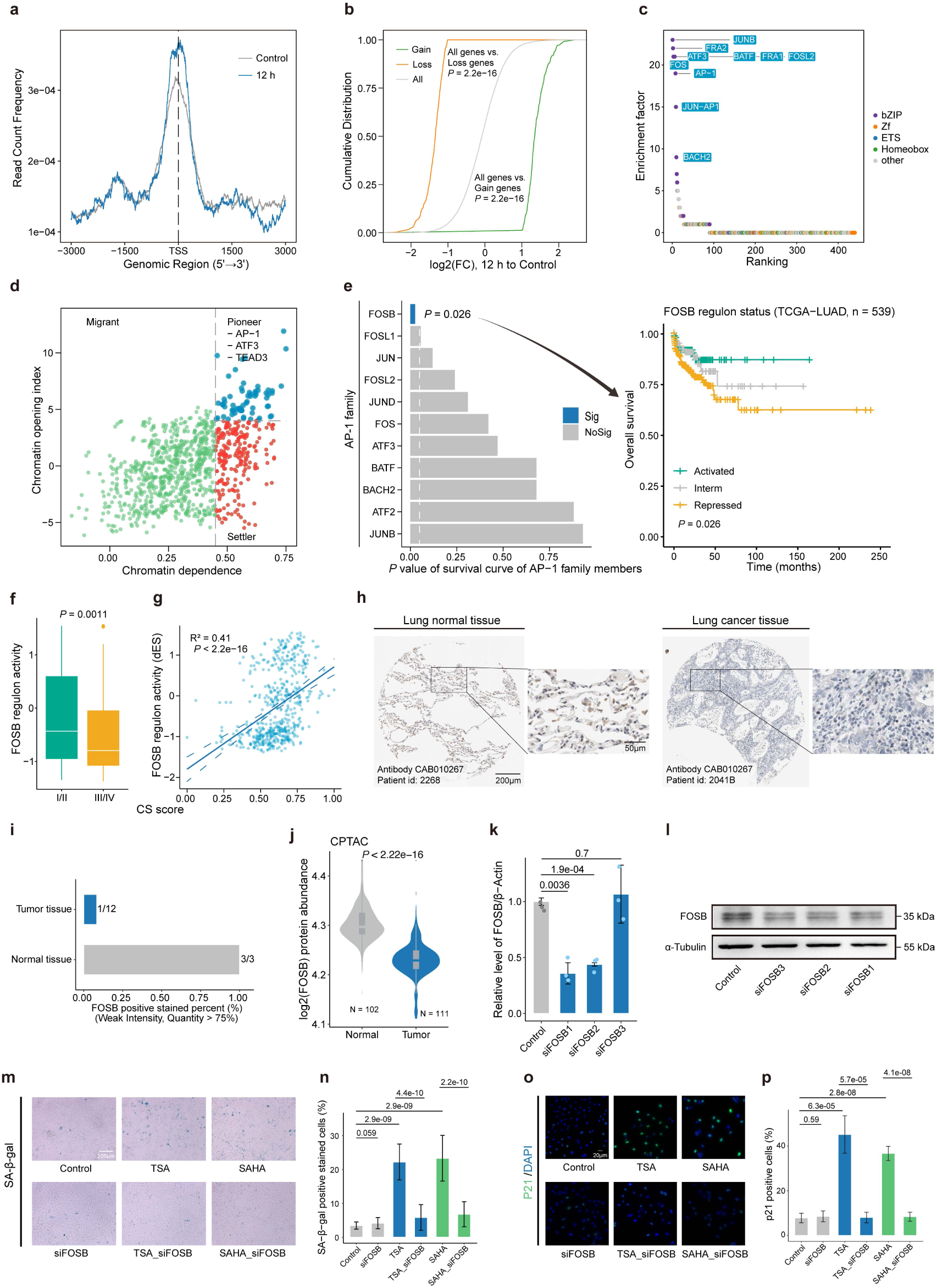
The AP-1 transcription factor (TF) contributes to HDACi-induced senescence. **a** Assay for transposase-accessible chromatin with sequencing (ATACseq) reads for A549 cells were compared between the control (gray) and trichostatin-a (TSA) 12-h treatment group (blue). **b** Cumulative distribution showing differences in chromatin accessibility between peaks related to cellular senescence (CS)-associated upregulated and downregulated genes (12 h vs. control). **c** Ranking of motifs enriched in senescence-associated open peaks (differential accessibility regions). Colors represent different TF families. **d** Categorization of TFs based on chromatin dependence (CD) and chromatin opening index (COI). Blue: pioneer factors with COI > 4 and CD > 0.45, Red: settler factors with COI < 4 and CD > 0.45, Green: migrator factors with CD < 0.45. The selected top pioneer factors are provided. **e** Comparison of *P* values for TF activity in the AP-1 family correlated with overall survival (OS) from survival analysis in The Cancer Genome Atlas (TCGA)-lung adenocarcinoma (LUAD) cohort (left). Blue: statistical significance. Kaplan–Meier survival curves showing the survival rate of FOSB regulon phenotypes with OS in the TCGA-LUAD cohort (right). **f** Box plots illustrating FOSB regulon activity across different American Joint Committee on Cancer (AJCC) stages in patients with LUAD in TCGA. The *P* values obtained from the Wilcoxon test are reported for comparisons between the two groups. **g** Scatter plots showing the correlation between FOSB regulon activity and the CS score in TCGA-LUAD (rank-based Spearman’s correlation). **h** Representative images of immunohistochemical staining for FOSB in normal lung (left) and tumor (right) tissues from the Human Protein Atlas (HPA). **i** Proportion of immunohistochemically positive staining of FOSB with weak intensity and quantity > 75% in tumor and normal tissues. **j** Violin plots showing the FOSB protein abundance in normal and tumor tissues of LUAD from Clinical Proteomic Tumor Analysis Consortium (CPTAC) proteogenomics data. **k** RT-qPCR analysis of FOSB expression following siRNA knockdown (*n* = 3). **l** Immunoblots showing the change in FOSB protein expression following knockdown. **m-p** Representative images and quantitative analysis of senescence-associated β-galactosidase (SA-β-gal) staining (m-n) and p21 immunofluorescence staining (o-p) in A549 cells subjected to different treatments, including control, siFOSB, trichostatin-a (TSA; 300 nM), vorinostat (SAHA; 500 nM), TSA with siFOSB, and SAHA with siFOSB, for 12 h. *n* = 6 biologically independent samples, with three randomly selected regions per sample analyzed. Scale bar: 200 μm for SA-β-gal staining and 20 μm for p21 immunofluorescence staining (Student’s *t*-test), and only the groups with significant differences are shown. Data are presented as the mean ± SD.

Potential regulatory transcription factors (TFs) were inferred using DARs and DEGs to identify CS-related TFs. Notably, TFs inferred from DEGs showed distinct TFs in various time points, with the unique TFs at 12 h belonging to the AP-1 family, including FOS, JUN, JUNB, FOSL1, and FOSL2 (Fig. 6f). Notably, the top ten enriched TFs inferred from DARs, which include JUNB, FRA2, ATF3, FOSL2, FOS, and AP-1, also belonged to the AP-1 family (Fig. 7c). Additionally, pioneer TFs may be crucial in controlling the senescence program by binding to compact chromatin areas, opening them, and recruiting other TFs to bind in their vicinity^44^. We identified pioneer factors by employing protein interaction quantitation (PIQ) based on ATAC-seq to calculate chromatin dependence (CD) and chromatin opening index (COI). AP-1 was identified as a pioneer factor that may bind to new peaks during HDACi treatment (Fig. 7d). In brief, transcriptomic and epigenetic analysis consistently revealed that the AP-1 family enhanced regulatory activity during HDACi-induced senescence.

### *FOSB*-mediated senescence reprogramming is critical for lung cancer

To assess the relationship between the regulatory activity of TFs within the AP-1 family and survival status in TCGA-LUAD patients, we performed regulon activity analysis for 11 members of the AP-1 family. Survival analysis revealed that only FOSB regulon phenotypes were significantly correlated with OS among 11 candidate regulators of the AP-1 family, and that a significantly better OS was observed under the activated FOSB regulon phenotype (*P* = 0.026; Fig. 7e). Meanwhile, regulon activity of FOSB was positively correlated with better clinicopathological characteristics of patients, including tumor pathological AJCC stage (Fig. 7f), as well as with CS score (Fig. 7g). Furthermore, immunohistochemical staining results from the Human Protein Atlas (HPA) showed that FOSB protein levels in lung tumor tissues were lower than in normal tissues (Fig. 7h, i). We also validated FOSB protein abundance in 213 LUAD patient samples using CPTAC proteogenomics data. FOSB levels were lower in LUAD tissues compared to normal tissues (*P* < 2.22e−16; Fig. 7j).

To further evaluate the function of FOSB in senescence, we performed *FOSB* knockdown in A549 cells via siRNA. Significantly decreased FOSB expression was observed through RT-qPCR and western blotting (Fig. 7k, l). Compared to the control, treatment with TSA or SAHA for 12 hours significantly increased SA-β-gal activity, p21 protein immunofluorescence, and TAF formation. In contrast, these senescence markers were markedly reduced following FOSB knockdown (Fig. 7m, n, o, p; Extended Data Fig. 5c, d). These results indicate that FOSB knockdown attenuates HDACi-induced senescence features and that FOSB could potentially serve as a novel driver molecule of senescence implicated in the onset and progression of LUAD.

## Discussion

The relationships between senescence levels and clinical characteristics, as well as the underlying mechanisms driving senescence programs in cancer, remain insufficiently understood. To address this gap, we employed several strategies. First, we developed a computational metric to quantify senescence levels. Second, we identified associations between senescence levels, molecular traits, and clinical outcomes. Third, we screened for drugs capable of inducing tumor senescence with potential antitumor effects. Finally, we elucidated key mechanisms underlying induced senescence in cancer and uncovered critical regulatory factors driving senescent cell reprogramming. Collectively, our computational framework enables the systematic evaluation of senescence levels in tumors, providing novel insights into their clinical relevance, therapeutic potential, and regulatory underpinnings across large-scale cancer cohorts.

Detecting senescence remains challenging due to phenotypic heterogeneity and genetic mutations in cancer^1,12^. To address this complexity, we developed PreCSenM, a computational model designed to quantify senescence levels through the following steps. First, we identified a robust senescence-related gene signature in non-cancerous cell lines and validated this consensus senescence gene signature (CSGS) in cancer datasets. The CSGS was functionally enriched in key senescence-related pathways and demonstrated superior performance in both normal and cancer cells compared to conventional signatures^14,18,20–24,30,31^. These findings suggest that CSGS may overcome current challenges posed by tumor heterogeneity and facilitate a context-independent understanding of CS. Second, we trained ten machine learning models using a leave-one-out cross-validation (LOOCV) framework based on the CSGS. This integrative approach enabled accurate evaluation of senescence-associated features across diverse biological and clinical contexts. We found that PreCSenM demonstrated greater robustness than existing detection methods^14–17,45,46^, and its performance was not confounded by differences in data types, cancer types, or cell types. In contrast, SenCID’s six overlapping submodels lack clear usage criteria, complicating selection and reducing reliability. Similarly, hUSI requires context-dependent submodule decisions, often leading to confusion and inconsistent outcomes. Third, we developed a user-friendly, web-based platform (http://precsenm.bmicc.org/) to make PreCSenM— a universal, context-independent model—readily accessible, in contrast to existing methods that require complex, disease-specific tuning. Unlike existing tools that require complex, disease-specific tuning, this platform supports integrated, multi-dimensional analysis, greatly enhancing the translational utility of CS-related discoveries. Nonetheless, our model has several limitations. Although CSGS improves model robustness, potential multicollinearity may obscure the contribution of individual genes. Additionally, when applied to bulk transcriptomic data, PreCSenM estimates the average senescence level across all cells in a sample. Further investigations are warranted to enable higher-resolution assessment of CS states within bulk transcriptomes, potentially by deconvolving cellular heterogeneity or integrating complementary single-cell datasets. Furthermore, as the morphological features are significant criteria of senescent cells, integrating the multiple transcriptomic markers with morphological features is needed to further improve the identification of senescent levels in the future.

We revealed the significant clinical value of the CS score in LUAD. As a leading cause of cancer-related death worldwide^32^, LUAD exhibits a strong association with cellular senescence, making it a compelling context in which to explore the complex role of senescence in cancer^47,48^. By integrating data from multiple LUAD cohorts along with diverse clinical parameters, our study demonstrated that the PreCSenM-derived CS score—unlike senescence estimates from other tools—showed significant associations with clinical outcomes. To uncover the biological mechanisms underlying the relationship between CS score and clinical features, we performed multi-omics analyses to characterize the molecular landscape of CS in LUAD. We found that lower CS levels in LUAD patients were associated with a progressive loss of genomic and epigenomic stability, along with an enhanced immune response. These findings underscore the CS score as a potential independent biomarker for prognosis and disease staging in LUAD, with multi-omic alterations in high-senescence patients suggesting a role for senescence in restraining tumor progression.

By CS score-based drug repositioning, we identified epigenetic drugs, including two HDACis (TSA and SAHA), as potential pro-senescence therapies that may improve outcomes in LUAD patients. Among two HDACis, the SAHA has received FDA approval for treating cutaneous T-cell lymphoma^49^. HDACis can induce multiple antitumor effects, including cell cycle arrest, senescence, and differentiation^50^. Our study discovered that HDACis can induce senescence in LUAD, with their underlying epigenetic mechanisms elucidated. Our results indicate that LUAD cells (A549 cells) treated with small-molecule HDACi displayed a senescent phenotype and decreased cell activity. Time-course analyses of transcriptomic and epigenomic data revealed that AP-1 family transcription factors (TFs) can reshape the epigenetic landscape during senescence. Notably, we identified FOSB, a relatively understudied AP-1 member, as a key regulator of senescence reprogramming in cancer cells and a potential predictor of improved survival in LUAD patients. These findings suggest that FOSB may serve as a promising therapeutic target for modulating senescence through epigenetic mechanisms. Previous studies have shown that HDAC inhibitors (HDACis) enhance the transcriptional activity of AP-1 by increasing histone acetylation^51,52^, and other reports suggest that HDACis may also regulate FOSB expression indirectly through additional signaling pathways^53^. Further investigations are needed to elucidate how HDACi-mediated activation of FOSB drives senescence in LUAD.

Overall, we have established a computational measure of senescence and identified associations between senescence and molecular characteristics, providing a tool available on GitHub (https://github.com/Lifei-Ma/PreCSenM) to evaluate senescent levels in various cancer types. Furthermore, we revealed that FOSB of the AP-1 superfamily is a key regulator for senescent signals and a potential diagnostic biomarker and therapeutic target for patients with LUAD. Hence, our work provides a framework for senescent evaluation and sheds new light on the development of future CS-related therapies.

## Methods

### Curation of CS cell line experiments

Our method depended on a sufficiently large number of available CS experiments well-established in the literature. For dataset selection, the following requirements had to be satisfied: (1) inclusion of datasets from RNA-seq and microarrays; (2) a single-channel array and not a custom-made one, allowing the use of standard annotations; (3) complete record of CS events in datasets; (4) at least two control and paired treatment datasets to provide either raw or processed data; (5) processing of the data using available packages; and (6) If the number of genes in the dataset is fewer than 16,000, it was not used as training data.

A comprehensive search was conducted for transcriptomic data related to CS in cell lines or tissues. A total of 888 cell lines from 68 publicly available datasets were accessed through the Gene Expression Omnibus (GEO) database and the ArrayExpress database at EMBL-EBI. Among these, 519 samples represent normal cell types, including Lung Fibroblasts, Foreskin Fibroblasts, Dermal Fibroblasts, Lung Epithelial, Mesenchymal Stem Cells, Umbilical Endothelial Cells, Eye Epithelial Cells, Myoepithelial Cells, Keratinocytes, Prostate Stromal Cells, Vascular Smooth Muscle Cells, Neural System Cells, and others. Additionally, 369 samples were derived from over 10 cancer types, including Brain, Breast, Leukemia, Liposarcoma, Liver, Lung, Melanoma, Osteosarcoma, Pancreatic, and Colorectal cancers. These selected datasets encompass various senescence-inducing conditions such as replicative senescence, DNA damage, gene editing, oncogene, oxidation, and others. Detailed information about these cell lines is provided in Extended Data Fig. 1a,b and Supplementary Tables 1 and 2.

### Publicly available multi-omics data and LUAD resources

RNA sequencing (Level 3 RNA-seq v2) data (transcripts per million, TPM), somatic mutation data (mutation annotation format), DNA methylation data (Illumina HumanMethylation 450 K-array), and associated clinical information from TCGA under TCGA-LUAD project were downloaded using the R package TCGAbiolinks (version 2.18.0)^54^. Nine independent public transcriptome datasets and complete clinical data for LUAD were retrieved from GEO, including GSE30219, GSE41271, GSE11969, GSE42127, GSE68465, GSE101929, GSE3141, GSE31210, and GSE37745. The GSE117570 dataset was used to determine the CS index using RNA-seq data for LUAD at the single-cell level.

### Cell culture

Human LUAD epithelial A549 cells were purchased from the American Type Culture Collection (Manassas, VA, USA) and cultured in RPMI-1640 medium (Gibco, Grand Island, USA) supplemented with 10% fetal bovine serum (Gibco), 100 U/mL penicillin (Gibco), and 100 μg/mL streptomycin (Gibco) in a humidified 5% CO_2_ incubator at 37 °C.

### Cell treatments for RNA-seq and ATAC-seq experiments

TSA and SAHA were purchased from MedChemExpress (New Jersey, USA) and dissolved in dimethyl sulfoxide (DMSO; Solarbio, Beijing, China) to achieve a storage concentration of 10 mM. For both RNA-seq and ATAC-seq, A549 cells were plated in six-well plates at a density of 2 × 10^5^ cells/well and exposed to a growth medium containing TSA (300 nM) or SAHA (500 nM). For the control conditions, cells were maintained in a growth medium containing 0.0033% DMSO, the same concentration as TSA, or 0.005% DMSO, the same concentration as SAHA, and incubated for 48 h without any additional treatment. Cells were exposed to TSA or SAHA at the aforementioned concentrations for 6, 12, 24, and 48 h. Duplicate experiments for each condition were performed to obtain samples for RNA-seq and ATAC-seq.

### RNA-seq library preparation and sequencing

Total RNA was extracted from A549 cells using the TRIzol method (Invitrogen, Carlsbad, CA, USA) and treated with RNase-free DNase I (Takara) according to the manufacturer’s instructions. A cDNA library with paired-end reads was constructed and sequenced on a NovaSeq 6000 system.

### ATAC-seq library preparation and sequencing

ATAC-seq was performed as previously described^55^. After drug exposure for different durations, live cultured A549 cells were scraped and placed in cold 1× phosphate-buffered saline (PBS). Purified nuclei were isolated from 50,000 cells and used for Tn5 enzyme transposition reaction and open area fragmentation of chromatin using the TruePrep DNA Library Prep Kit V2 for Illumina (Vazyme Biotech Co., Ltd.) and incubated at 37 °C for 30 min. The DNA was purified using a kit (Qiagen), following the manufacturer’s instructions. DNA libraries were generated by performing nine cycles of PCR amplification using the NEBNext High-Fidelity 2 PCR Master Mix (New England Biolabs). After size selection to remove non-target fragments, the DNA libraries were sequenced on an Illumina NovaSeq 6000 platform, generating 150 bp paired-end reads for the ATAC-seq libraries.

### RNA-seq and microarray processing

RNA-seq raw sequencing data, in the form of fastq files, were downloaded from SRA (https://www.ncbi.nlm.nih.gov/sra) using prefetch and fastq-dump in the SRA Toolkit (version 2.11.0) or by sequencing samples using the NovaSeq 6000 sequencing system. Quality control was conducted using FastQC software (version v0.11.9), and low-quality reads with an average quality score < 20 were discarded. Adapter sequences and low-quality bases were trimmed using trim-galore (version 0.6.7). The aligned sample sequences were mapped to the GRCh38 genome using STAR (version 2.5.2a)^56^, and gene expression (TPM) was quantified using RSEM (version v1.3.3)^57^. Protein-coding genes were identified using the BiomaRt R package (version 2.46.3). After excluding non-coding genes, 19,425 protein-coding genes were included in the subsequent analyses. The normalized gene expression was log2 transformed. The DESeq2 package (version 1.30.1) in R was used to compare DEGs between the treatment and control groups for each condition. GO analyses were performed using the ClusterProfiler (version 3.18.1) package in R to investigate the potential biological functions of DEGs. Raw microarray data were retrieved from the GEO database, which was generated using various platforms, including Affymetrix GPL570 (Human Genome U133 Plus 2.0 Array), GPL96 (Human Genome U133A Array), GPL6244 (Human Gene 1.0 ST Array), GPL5175 (Human Exon 1.0 ST Array), GPL7015 and GPL6884 (HumanWG-6 v3.0 expression beadchip), Illumina GPL10558 (HumanHT-12 V4.0 expression beadchip), Agilent GPL6480 (Whole Human Genome Microarray 4×44K G4112F), and Agilent GPL13497 (Whole Human Genome Microarray 4×44K v2). Raw data from the Affymetrix platform underwent quality control and normalization using the Robust Multichip Average method implemented in the oligo package (version 1.54.1)^58^. The Illumina and Agilent platforms were processed using the limma package (version 3.46.0). The annotation package available or the GPL platform files of the dataset were used to match the probe annotation with HUGO Gene Nomenclature Committee (HGNC) symbols. Probes that did not match the HGNC symbols were eliminated, and the average intensity of the probes matching the same HGNC symbols was computed. All samples for training were merged for the machine learning procedure, and batch effects were eliminated using the ComBat algorithm in the sva package (version 3.38.0)^59^. Gene expression was rescaled via z-score transformation across all samples.

### ATAC-seq processing

Trim_galore (version 0.6.7) was used to remove poor-quality reads and adapter sequences of raw sequence reads. Clean data were aligned to the human reference genome (hg38) using Bowtie2 (version 2.3.4.2). Uniquely mapped paired reads were retained, and SAM files were sorted and converted to the BAM format using Samtools (version 1.3.1). Subsequently, peak calling was performed using MACS2 (version 2.2.7.1) to identify the significant peaks in each sample. Peak regions of the genome were annotated using the ChIPseeker R package (version 1.26.2). The Integrative Genomics Viewer tool converted the viewable bigwig files from DeepTools (version 3.5.1). To identify DARs, the treatment and control groups were compared in separate cell lines using DESeq2 (version 1.30.1) with a threshold of *P* < 0.05.

### Generated signatures, model building, and validation using machine learning-based ensemble methods

To develop a consensus CS signature and prediction model, 11 machine learning algorithms were incorporated to enhance the accuracy and stability of the results. These algorithms included Boruta, LMM, LR, Ridge, Lasso, Enet, RF, XGBoost, SVM, ANN, and PLSR. The senescence signature was generated, and model development and validation proceeded as follows. First, approximately 70% of the normal senescence cell lines, selected based on the dataset selection criteria, were randomly allocated for training to identify feature genes and construct a robust prediction model, with this process repeated 10 times. The remaining 30% of normal senescence cell lines were used as the testing set, while tumor cell lines or tissues were utilized as the validation set. Second, the Boruta feature selection algorithm, an ensemble learning model for feature selection, was applied to the training dataset to identify a refined consensus CSGS. Third, functional enrichment analysis of genes associated with senescence signatures was conducted using Metascape^60^. Following this, ten binary classification algorithms (excluding Boruta) were performed on the senescence signatures generated in the previous step to fit prediction models, with the best hyperparameters identified based on the LOOCV framework in the training set. All models were tested on the remaining testing set and tumor datasets, exhibiting distinct senescence induction patterns. To ensure robustness, model evaluations were repeated at least ten times with randomly partitioned training and testing datasets. Both microarray and RNA-seq data were used in training, testing, and validation sets. Finally, the accuracy, precision, recall, F1 score, AUC of the receiver operating characteristic (ROC) curve, and area under the precision-recall curve (AUPRC) were evaluated for each model across all training, testing, and tumor datasets. After weighing all metrics based on sample size, the model with the highest weighted metrics was considered optimal.

### Model description

One feature-based models and ten binary classifiers were implemented, totaling 11 models, using the R packages, including lmerTest (version 3.1.3), glmnet (version 4.1.2), caret (version 6.0.88), randomForest (version 4.6.14), Boruta (version 7.0.0), xgboost (version 1.5.0.2), e1071 (version 1.7.6), neuralnet (version 1.44.2), and pls (version 2.8.0). The Boruta feature selection algorithms was employed to identify the crucial genes associated with CS. We utilized Boruta, a feature selection algorithm designed specifically for ensemble learning models, to enhance the precision of the results. The response variable was senescent status, with 0 representing non-senescent and 1 representing senescent status. This approach generated a compact and high-quality set of senescence-related feature genes for further analysis. Predictive binary classification models were then constructed using the significant senescence-related signature genes as input variables.

The LOOCV scheme was implemented to identify the optimal hyperparameters in the training set for each model to build the best-performing classifier. A dataset was divided into *n* portions, where *n* is the number of samples in the training set. In each iteration, n-1 portions were used as the training set to develop the model, and the remaining portion was designated the validation set for model evaluation (n = training sample number). This process was repeated *n* times until each portion was used as validation data once. For predictive models, the best signature genes were used as input variables in the Ridge, Lasso, Enet, RF, XGBoost, SVM, ANN, and PLSR algorithms. In contrast, each gene was used individually as an input variable in the LMM and LR models. The senescence scores for the LMM and LR models were calculated by multiplying the gene expression of the best signature genes (Ei) in each sample with their respective coefficient (βi) as follows:

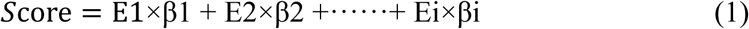

Data were scaled to have the same mean of 0 and a standard deviation of 1, and min–max normalization was performed to map the CS probability score of each sample to the 0–1 range to account for the differences in the strength of gene expression signatures and enable simultaneous comparison of relative scores across samples. A higher score indicates a higher probability of experiencing a senescent status, where the default cut-off is 0.5, the median of scores, or adjusted based on specific situations.

### Evaluation of model performance

Model performance was evaluated using the accuracy, precision, recall, F1 score, AUC of the ROC curve, and AUPRC. All metrics and plots were generated using the ROCR package (version 1.0.11) or a confusion matrix. Accuracy is the ratio of true positive and true negative predictions to all predictions and is calculated as follows:

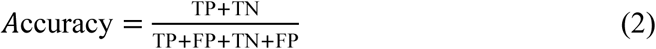

where FP represents the number of false positive cases, TP represents the number of true positive cases, and TN represents the number of true negative cases. Precision is the ratio of positive examples that are truly positive and was calculated as follows:

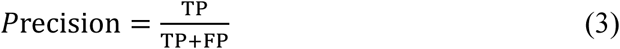

Recall is the sensitivity of predictions and is calculated as follows:

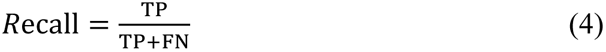

where FN represents the number of false negative cases.

The F1 score combines precision and recall into a single number and was calculated as follows:

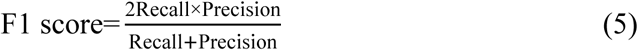

The ROC curve plots of the true-positive rate against the false-positive rate for all possible thresholds. The AUPRC curve, a measure of a model’s predictive performance, is based on the relationship between the positive predictive value (precision) and true positive rate (recall) for detecting samples.

### Comparison to public senescent signatures and senescence gene scoring algorithms

To evaluate the performance of PreCSenM in predicting senescent cells, we compared it against single canonical markers (CDKN1A (p21) and CDKN2A (p16))^30,31^, five publicly available senescence signatures^14,18–24^, and four senescence gene scoring algorithms^14–17^ (Supplementary Table 3) across the training, testing, and tumor datasets. All algorithms were implemented following the protocols described in their original publications. The performance of each signature was assessed by measuring the AUC-ROC, as described previously.

### Validation of CS scores in various datasets

The scores were measured using the PreCSenM method to verify the suitability of the CS scores for various conditions and data types. Initially, to evaluate the suitability of PreCSenM for analyzing scRNA-seq data, we used publicly available single-cell transcriptomics datasets, with the senescent status reported (GSE115301). Additionally, we selected TCGA data containing normal and cancer samples from ten cancer types with more than 35 samples as previously reported. To assess the statistical differences in CS scores between distinct pairs, a two-sided Mann–Whitney U-test (Wilcoxon test) was conducted for non-normally distributed data, and the corresponding *P* values were determined.

### CS score versus clinical predictors

To elucidate relationships between the CS score, OS, and clinical characteristics of patients with LUAD, a survival analysis was performed on ten independent datasets, including the TCGA and GEO databases. The optimal cutoff for all samples divided into high- and low-CS groups was determined by the “surv_cutpoint” algorithm in the Survminer (version 0.4.9) R package. Next, Kaplan–Meier survival curves were used to assess differences in survival between these two groups and the log-rank test was used to identify significant differences (*P* < 0.05). Survival analyses were conducted using R survival (version 3.1.12) and Survminer packages. A comprehensive analysis investigated the correlation between the CS score and various clinical features, such as TNM and AJCC stage, and vital statuswithin the two distinct CS groups.

### GSVA

Pathway analyses were performed on 34 senescence-related pathways among the 50 hallmark pathways reported in the Molecular Signatures Database (MSigDB), exported using the GSEABase package (version 1.52.1). Pathway activity estimates for individual patients^45^ were derived using normalized enrichment scores (NES) from GSVA, performed with default parameters in the GSVA package (version 1.38.2).

### Genomic variation analysis

The SCNA score, which reflects the level of SCNAs of a tumor, was generated as previously described^33^. SCNA profiles were examined using segmented files (nocnv_grch38.seg) from TCGA as input files using GISTIC 2.0^61^ to identify the chromosome, arm, and focal levels in TCGA-LUAD patients. Alterations occurring in >70% of an arm were considered broad events. Significant peak regions were found with a q-value <0.25, and the confidence level was set at 0.95. The ratios of log2 copy number for focal-level events were scored as follows: 2 if log2 ratio ≥ 1, 1 if log2 ratio < 1 and ≥ 0.25, 0 if log2 ratio < 0.25 and ≥ −0.25, −1 if log2 ratio < −0.25 and ≥ −1, and −2 if log2 ratio < −1. The focal score of a tumor is the absolute sum of all focal-level scores in the tumor. Similar methods were used to define scores at the arm and chromosome levels. Chromosome-level events were defined as those in which the log2 ratios in both arms were identical. Total SCNA scores at the focal, arm, and chromosomal levels were used to determine the overall SCNA score for the tumor. The relationship between CS and SCNA scores for LUAD was examined using simple linear regression.

To determine the relationship between somatic variants and CS in LUAD, Mutation Annotation Format files were analyzed and visualized using the R package maftools (version 2.6.5) with standard process settings. Mutant genes were selected in accordance with those of classic senescence-related pathways. The variant classifications for each sample were uniformly summarized and displayed without discrimination. Mutations in various pathways were computed for distinct senescent subgroups.

TMB refers to the number of gene mutations in each patient, including base substitutions, insertions, and deletions. For each LUAD sample, TMB was computed as the sum of the nonsynonymous mutations divided by the number of exons (38 million). Associations between the CS and TMB scores were calculated using the same linear model as the SCNA score.

For each patient, the AS (available at the NCI Genomic Data Commons), another indicator of genome instability, was obtained as the sum of altered chromosome arms^62^. MSI was calculated using the PreMSIm algorithm as a genomic property of cancers with impaired DNA mismatch repair.

### DNA methylation analysis

Owing to the scale of the available DNA methylation platform, Illumina HumanMethylation 450k array, it was possible to identify the association between DNA methylation and CS in patients with LUAD using the R package ChAMP (version 2.20.1). DNA methylation indices were displayed as β values, representing the ratios of the intensities used to quantify DNA methylation levels. Thorough filtering was performed using the following criteria: probes with *P* values > 0.01 or with less than three beads in at least 5% of samples were removed, along with non-CpG probes, SNP-related probes, multi-hit probes, and probes located on the X and Y chromosomes^63,64^. DMPs with a mean β value > 0.2 or < 0.2, indicating an adjusted *P* value < 0.05, were identified as significant hypermethylators or hypomethylators, respectively, using the default parameters of the champ.DMP function in the ChAMP package. DNA methylation levels of DMPs in the two CS subgroups of LUAD were compared.

### Single-cell RNA-seq data analysis

Gene expression matrices for LUAD were obtained by extracting scRNA-seq data for lung tumors (GSE117570), which were then preprocessed using the standard analysis pipeline in MAESTRO^35^. To select a subset of high-quality cells, those with < 1000 unique reads or UMIs, < 500 expressed genes or > 5% UMIs from the mitochondrial genome were discarded; the remaining cells were used for downstream expression quantification and clustering analysis. Based on prior knowledge of marker genes, cell clusters from scRNA-seq were annotated with three distinct major cell types: immune, stromal, and malignant cells.

The CS score of scRNA-seq data was calculated by multiplying the gene expression and coefficient of 236 CSGSs in each cell. Different cell types and tumor T stages were assessed to determine their senescent states. The median score was used as the default cutoff for two distinct groups of malignant cells, where a higher score denotes a higher likelihood of senescence.

To identify senescence-associated DEGs between the high- and low-CS groups in malignant cells, the FindMarkers function in the R package Seurat (version 4.0.1)^65^ was used based on the Wilcoxon rank sum test. Genes were selected when the *P* value was < 0.05 and the average log2-transformed difference between the two groups was > 0.15. Biological pathway enrichment of DEGs from scRNA-seq data was analyzed using the Reactome database (https://reactome.org).

To investigate cell–cell communication networks via ligand–receptor interactions, a count matrix of four different cell types (immune, stromal, high-CS-malignant, and low-CS-malignant cells) was used as the input in CellPhoneDB (version 1.1.0). Based on the mean gene expression of the ligand from one cell type and the corresponding receptor from another, permutation tests between the two cell types were performed to identify potentially important interaction pairs. Pairs with average expression > 0.2 and *P* value < 0.05 were designated as significant predicted interaction pairs.

### Generation of query signatures for drug screening

Candidate genes were identified in the TCGA-LUAD dataset between the two CS groups. Hierarchical filtering was performed based on the number of senescence- and survival-related features between the two groups to reduce the list of candidate genes; only genes with more than 100 related features were retained for further research. The DEGs between the high- and low-CS groups were analyzed using the DESeq2 package in R. Adjusted *P* values were applied to correct false-positive results using the default Benjamini–Hochberg false discovery rate (FDR) approach. Threshold values of |log FC| > 2 and FDR < 0.05 were used to identify significant genes. A univariate Cox proportional hazard regression model was developed to assess the effects of various genes on OS using the DEGs obtained from the previous step. Genes with a cutoff value of *P* < 0.05 were defined as candidate genes related to OS and were selected for the next step. In the third feature selection stage, the RSF algorithm was performed using the randomForestSRC (version 2.14.0) R package to reduce the number of candidate DEGs further^66^. The RSF model was constructed according to the optimal parameters, and 1000 trees were grown using the log-rank splitting rule, which was repeated 1000 times. Finally, genes were screened according to the importance of positive values and considered optimal candidates.

### LINCS and cancer cell line–drug response data source and processing

The LINCS program has made more than 1.3 million transcriptomic profiles available using the L1000 platform^67^, which collects the gene expression profiles of several cell lines under the induction of various perturbagens. LINCS level 5 data (moderated Z-score) were obtained from the GEO database (Phase I: GSE92742, Phase II: GSE70138), which covers differential expression signatures for nearly 20,000 unique compounds and meta-information. This study used more than 25,446 A549 lung cancer cell lines and their corresponding data. In addition, the mechanisms of action (MoA) and clinical phase information for compounds in LINCS were obtained from the Drug Repurposing Hub (https://clue.io/repurposing)^68^. Response data with IC50 values were evaluated using the PRISM repurposing dataset (22Q4, released December 2022)^69^ to assess the drug sensitivity of repositioned candidates across lung cancer cell lines.

### Signature matching approach for drug prediction

Three distinct benchmark approaches, KS^36^, XSum^37^, and RGES^38^, were applied to perform signature matching analysis with the optimal query signature obtained from the previous step.

In this study, the KS approach divided query signatures into discrete positive and negative regulators. The maximum deviation (MD)-based enrichment scores (ES) of positive (ES_pos_) and negative (ES_neg_) regulators were estimated using all compound profiles as references. The KS score was defined as KSscore = ES_pos_ − ES_neg_, with the enrichment scores for the positive and negative regulators having the same sign, KSscore = 0. The KS approach is the most extensively used for signature matching, particularly in CMap studies^36^. Similar to KS, the XSum approach involves splitting query signatures into two gene sets. Briefly, the sum of the change values in the reference drug signatures relative to positive (Sum_pos_) and negative (Sum_neg_) regulators was calculated. The XSum score was calculated as follows: XSum = Sum_pos_ – Sum_neg_. The RGES approach is an improvement of the original KS method, which was recently shown to perform better in drug prediction^38^. RGES is a metric that illustrates the inverse correlation between the gene expression profiles of a given disease and its treatment. The RGES is defined as ES_pos_ − ES_neg_, regardless of the direction.

Rank aggregation analysis, an order statistics-based method proposed by Stuart et al.^39^, was conducted to obtain a robust order of drugs based on the ranking results of the methods mentioned above. Rank aggregation scores (RASs) represent the probabilities of being used to generate a final ranking list of potential drugs, and drugs with a higher RAS denote greater reversal potency in tumors. The RAS is defined as

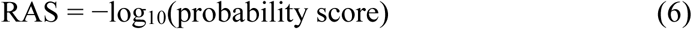

### TF enrichment analysis using RcisTarget

The potential regulatory TFs of DEGs from different comparisons were predicted using the R package RcisTarget (version 1.10.0) based on TF-binding motifs with default parameters. The TF networks were constructed using Cytoscape (version 3.9.1).

### Motif enrichment analysis

Motif enrichment analysis was performed based on the genomic regions of the DARs from the ATAC-seq data using the findMotifsGenome.pl function in HOMER (version 4.10).

### Identification of pioneer TFs using PIQ

Pioneer factors were identified using the PIQ algorithm with the published code as proposed by Sherwood et al.^70^ based on ATAC-seq datasets. The PIQ allows for the functional categorization of TFs into settlers, migrants, and pioneers. Pioneer factors were defined as TFs with CD > 0.45, COI > 4; settler factors as TFs with CD > 0.45, COI < 4; and migrator factors as TFs with CD < 0.45.

### Regulon activity analysis

The relative activities of 11 candidate regulator genes previously reported to belong to the AP-1 family were determined^71^. The “regulator” is a gene whose product stimulates and/or represses the target set of genes, namely the “regulon“^72^. Transcriptional regulatory networks (regulons) for the inferred AP-1 family-associated TFs were reconstructed using the R package RTN (version 2.14.1)^72^ based on TCGA-LUAD RNA-seq data. Specifically, mutual information and Spearman’s correlation indicated potential associations between a regulator and all targets in the transcriptome expression profile. A regulatory network was constructed by removing associations in accordance with permutation analysis (FDR < 1e-5), and unstable interactions were eliminated using bootstrapping (*n* = 1000 resamples, consensus bootstrap > 95%). Indirect interactions were eliminated by applying the Data Processing Inequality algorithm. Using the two-tailed GSEA method, an enrichment score (dES) was assigned to each individual in a cohort. The cohort samples were stratified into three categories according to the ranking of the dES values, as follows: activated, intermediate, and repressed. Based on these classifications, survival-associated regulators were identified using survival analyses, which showed the prognostic value of impressive regulon phenotypes for OS outcomes.

### RNA interference

Two duplex RNAi oligos (Synbio Technologies) targeting human FOSB mRNA sequences were synthesized to knockdown endogenous gene expression (Supplementary Table 5). siRNAs were transfected using the RNAiMax Reagent (Invitrogen) according to the manufacturer’s instructions at a final concentration of 50 nmol/L. The efficiency of siRNA transfection was assessed using immunoblot analysis and RT-qPCR.

### RNA collection and RT-qPCR

RNA was extracted from treated A549 cells using TRIzol reagent, and cDNA was synthesized using PrimeScript™ IV 1st strand cDNA Synthesis Mix (Takara, China). RT-qPCR was performed in triplicate using Taq Pro Universal SYBR qPCR Master Mix (Vazyme Biotech Co., Ltd.) on a Light Cycler 480 instrument (Roche). Gene expression was quantified using the ΔΔCT method and normalized to the expression level of the β-actin gene. The primer sequences for RT-qPCR were as follows: *FOSB* forward, 5′– GCTGCAAGATCCCCTACGAAG–3′; *FOSB* reverse, 5′–ACGAAGAAGTGTACGAAGGGTT–3′; β-actin forward, 5′– GGGAAATCGTGCGTGACATTAAG–3′; and β-actin reverse, 5′– TGTGTTGGCGTACAGGTCTTTG–3′.

### Western blotting

Treated cells were lysed using RIPA buffer containing protease inhibitor cocktail (Thermo Fisher, MA, USA; 78430). Approximately 40 μg of total protein was separated by sodium dodecyl sulfate-polyacrylamide gel electrophoresis and transferred onto polyvinylidene fluoride membranes. The membranes were then blocked using 5% non-fat milk before incubating overnight at 4 °C with antibodies against FOSB (1:10000, ab184938, Abcam) and alpha-tubulin (1:2000, t8203, Sigma). The membranes were blotted with rabbit monoclonal antibodies at room temperature for 1 h, and the bands were detected using an enhanced chemiluminescent reagent (Thermo Fisher).

### Cell viability

Cells were cultured until confluent in 96-well plates and then incubated with TSA or SAHA at the optimal concentrations (300 or 500 nM, respectively) for 96 h. Cell viability was assessed using CCK-8 cell proliferation and cytotoxicity assay kits (Solarbio, CA1210), and absorbance at 450 nm was measured using a microplate reader (Thermo Fisher).

### SA-β-gal assay

The SA-β-gal activity was determined using a senescence detection kit according to the manufacturer’s instructions (Beyotime). A549 cells were washed twice with 1× PBS and fixed with fixing buffer for 15 min at room temperature. The cells were washed thrice with PBS and stained with staining mixture at 37 °C overnight. The number of SA-β-gal^+^ senescent cells was counted at five different fields in each well under a microscope, and the proportion of SA-β-gal^+^ cells to total cells was calculated using ImageJ software (NIH).

### Immunofluorescence staining

A549 cells were washed three times with PBS, fixed with 4% paraformaldehyde for 20 min, and permeabilized with 0.1% Triton X-100 for 15 min at room temperature. Then, samples were blocked with 5% bovine serum albumin (BSA) for 40 min at room temperature and stained with a rabbit monoclonal antibody against p21 (1:1000, Abcam, ab109520) diluted in PBS containing 1% BSA overnight at 4 °C. Alexa Fluor 488 goat anti-rabbit IgG (Abcam, ab150077) was used as the secondary antibody. Finally, the samples were prepared for DAPI nuclear staining and observed under a fluorescent microscope.

### TAF Staining and Quantification

For A549 cells, sections were fixed with 4% paraformaldehyde (Servicebio) for 15 min, followed by three washes with PBS and permeabilization with 0.3% Triton X-100 in PBS for 15 min. After three additional PBS washes, samples were denatured in a hybridization buffer containing 70% formamide and 2× SSC at 85°C for 10 min. The telomere-specific probe (Servicebio) was pre-denatured at 85°C for 5 min prior to use. Sections were then washed three times with 2× SSC and incubated overnight at 40°C with the denatured telomere probe diluted to 300 nM in hybridization solution containing 35% formamide, 2× SSC, 10 mM Tris-HCl (pH 7.2), and 100 μg/mL salmon sperm DNA (Thermo Fisher). Subsequent hybridization steps were performed sequentially with branch probes and signal probes at 40°C for 2 h each.

For γ-H2AX immunofluorescence, sections were processed using a DNA Damage Assay Kit (Beyotime) according to the manufacturer’s instructions. Briefly, cells were incubated with primary anti-γ-H2AX antibody at 4°C overnight, followed by incubation with a fluorophore-conjugated secondary antibody at room temperature for 1 h. Samples were washed three times with PBS and mounted using ProLong™ Gold Antifade Mountant with DAPI (Invitrogen). All incubations and washes were performed under controlled humidity and temperature conditions to ensure optimal signal retention.

### Statistical analysis

Statistical analyses and graphical visualizations were performed using R software (version 4.0.2). The Wilcoxon test compared two groups with non-normally distributed data, whereas the Student’s *t*-test was used for normally distributed data. Two-sided Kruskal–Wallis tests compared more than two groups with non-normally distributed data. Fisher’s exact test was used to analyze contingency table variables. Correlations between two continuous variables were measured using Spearman’s correlation. The statistical significance of the survival analysis was determined using a log-rank test. Data are presented as the mean ± standard deviation (SD) for cell-line experiments. Statistical information for all experiments is detailed in the respective figure legends.

## Acknowledgements

This work was supported by grants from National Natural Science Foundation of China (82030017, 82225007 and 82073082), Chinese Academy of Medical Sciences Innovation Fund for Medical Sciences (CIFMS2021-I2M-1-050 and CIFMS2022-I2M-2-002), and National High Level Hospital Clinical Research Funding (2022-PUMCH-C-008 and 2022-PUMCH-C-014). This study was also supported by the Medical Science Data Center at Shanghai Medical College of Fudan University and the High-performance Computing Platform of Suzhou Institute of Systems Medicine, Chinese Academy of Medical Sciences & Peking Union Medical College.

## Author contributions

L.M., H.L., and H.-Z.C. conceived and designed the study. L.M., H.L., and H.-Z.C. planned and designed the experiments, interpreted the data and wrote the manuscript. L.M. performed computational analyses, designed bioinformatics pipelines and prepared figures. H.L. generated the cell culture systems and performed the RNA-seq and ATAC-seq experiments. All authors participated in data discussion and manuscript reviewing and approved the submitted manuscript.

## Data availability

All high-throughput sequencing data generated in this study have been deposited in the GEO database under the accession numbers GSE244427 and GSE245231. The previously published CS experiment data analyzed in this study are available under the accession numbers provided in Supplementary Table 1. Tumor CS experiments data analyzed in this study are available under the accession numbers provided in Supplementary Table 2. The scRNA-seq validation data with the senescent status can be accessed at GEO under the accession number GSE115301, and the LUAD scRNA-seq data is available under the accession number GSE117570. The protein abundance data for LUAD is available in the Clinical Proteomic Tumor Analysis Consortium (CPTAC) proteogenomics portal (https://proteomics.cancer.gov/data-portal/). All experimental materials generated in this study are available from the corresponding author upon request. Immunohistochemical staining images were downloaded from HPA (https://www.proteinatlas.org/).

## Code availability

The source code used to calculate the CS score of PreCSenM and other analyses is freely available at https://github.com/Lifei-Ma/PreCSenM.

## Extended data

**Extended Data Fig. 1.**
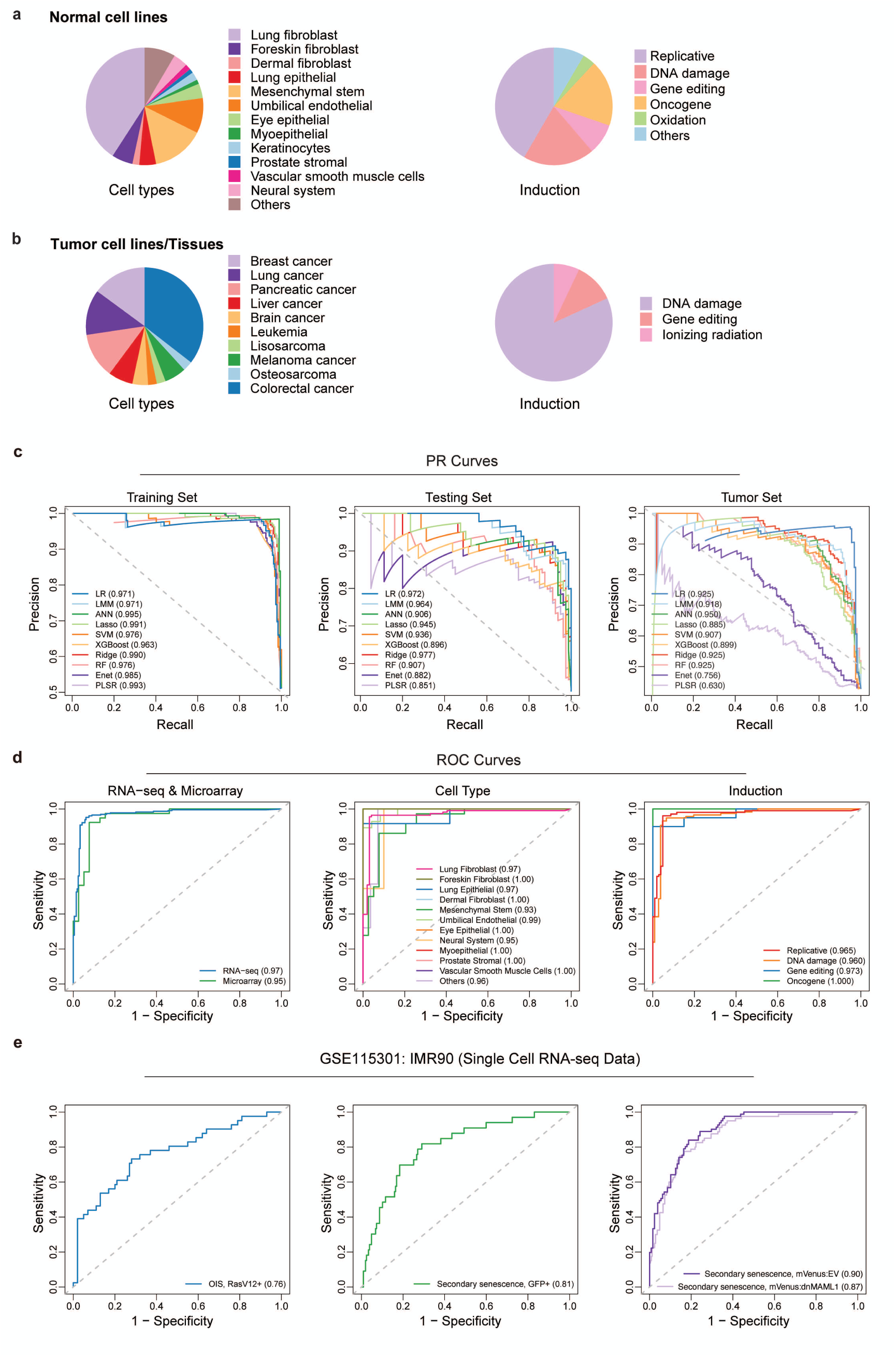
Cellular senescence data composition and model performance assessment for evaluating senescence levels. **a-b** The composition of cell types and senescence-inducing treatments in normal cell lines (a) and tumor cell lines or samples (b). **c** Corresponding area under the precision-recall curve (AUPRC) values of ten machine learning models are shown in the training (left), testing (middle) and tumor (right) sets. **d** Receiver operating characteristic (ROC) curves and area under the curve (AUC) values were evaluated for normal senescent cell types across three classifications: data type (left), cell type (middle), and induction type (right). The AUC represents the average true positive rate (sensitivity) across different false positive rate (1 − specificity) thresholds. **e** Receiver operating characteristic (ROC) curves and corresponding area under the curve (AUC) values for PreCSenM were assessed using IMR90 single-cell RNA-seq data (GSE115301). The analysis included three datasets: (left, *n* = 141) oncogene-induced senescence in RasV12-positive cells, (middle, *n* = 240) secondary senescence in GFP-positive IMR90:GFP cells, and (right, *n* = 536) secondary senescence induced by compromised Notch signaling, where IMR90 cells expressing a dominant-negative form of mastermind-like protein 1 (mVenus:dnMAML1) or an empty vector control (mVenus:EV) were co-cultured with ER:Ras IMR90 cells in the presence of tamoxifen. The AUC value represents the average true positive rate (sensitivity) across different false positive rate (1 − specificity) thresholds.

**Extended Data Fig. 2.**
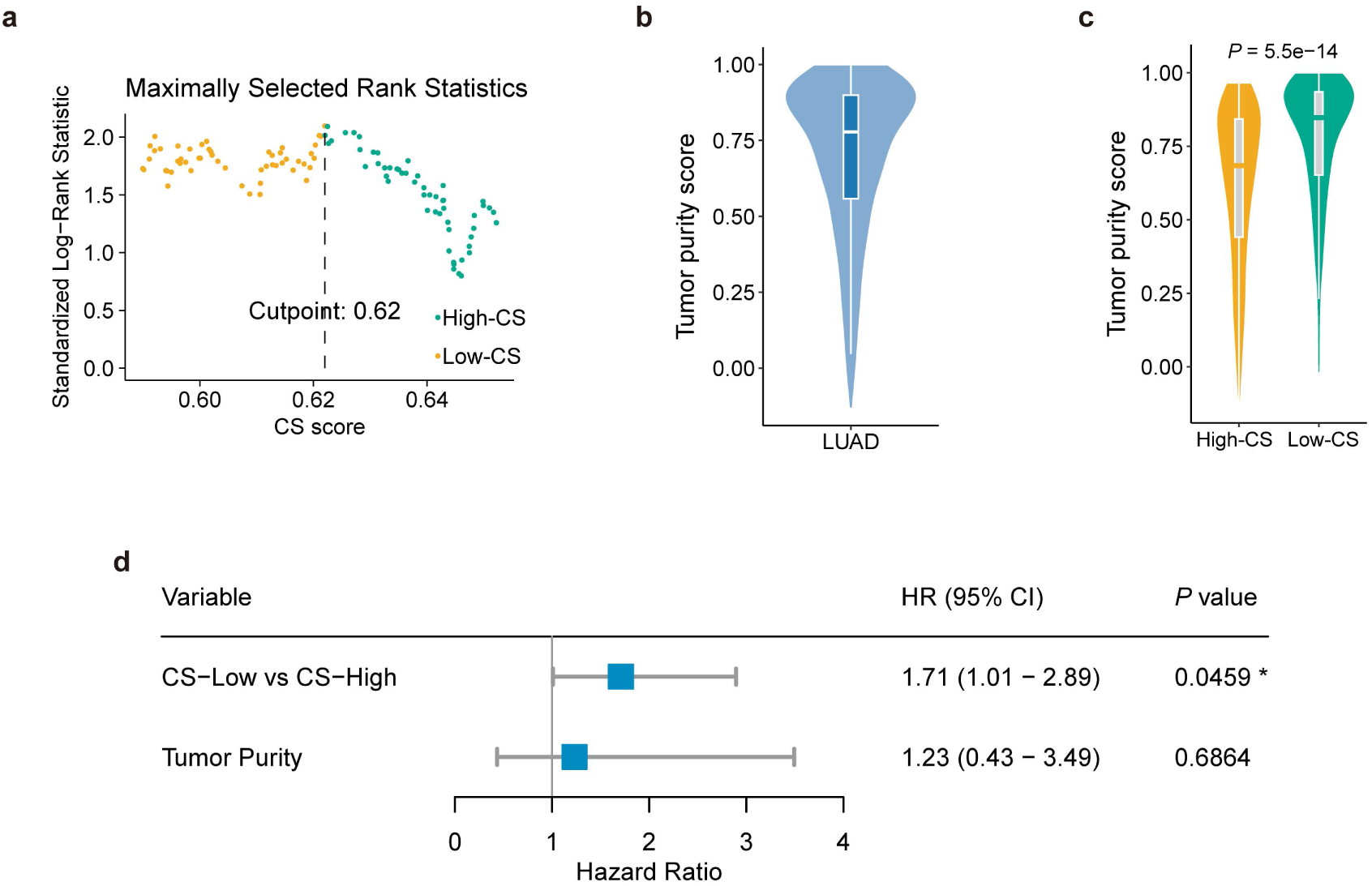
Stratification of TCGA LUAD patients based on senescence groups and assessment of tumor purity. **a** Scatter plot of the optimally selected rank statistic indicates that a cut-off value of 0.62 was ideal for dividing patients with LUAD into high- and low-CS groups (CS high vs. CS low: *n* = 240 vs. 299). **b** Violin plot showing the distribution of tumor purity scores in TCGA LUAD. Tumor purity was assessed using the ESTIMATE algorithm. **c** Violin plot showing the distribution of tumor purity scores between the two CS subgroups in TCGA LUAD. **d** Cox regression analysis of overall survival in TCGA-LUAD CS subgroups, adjusted for tumor purity.

**Extended Data Fig. 3.**
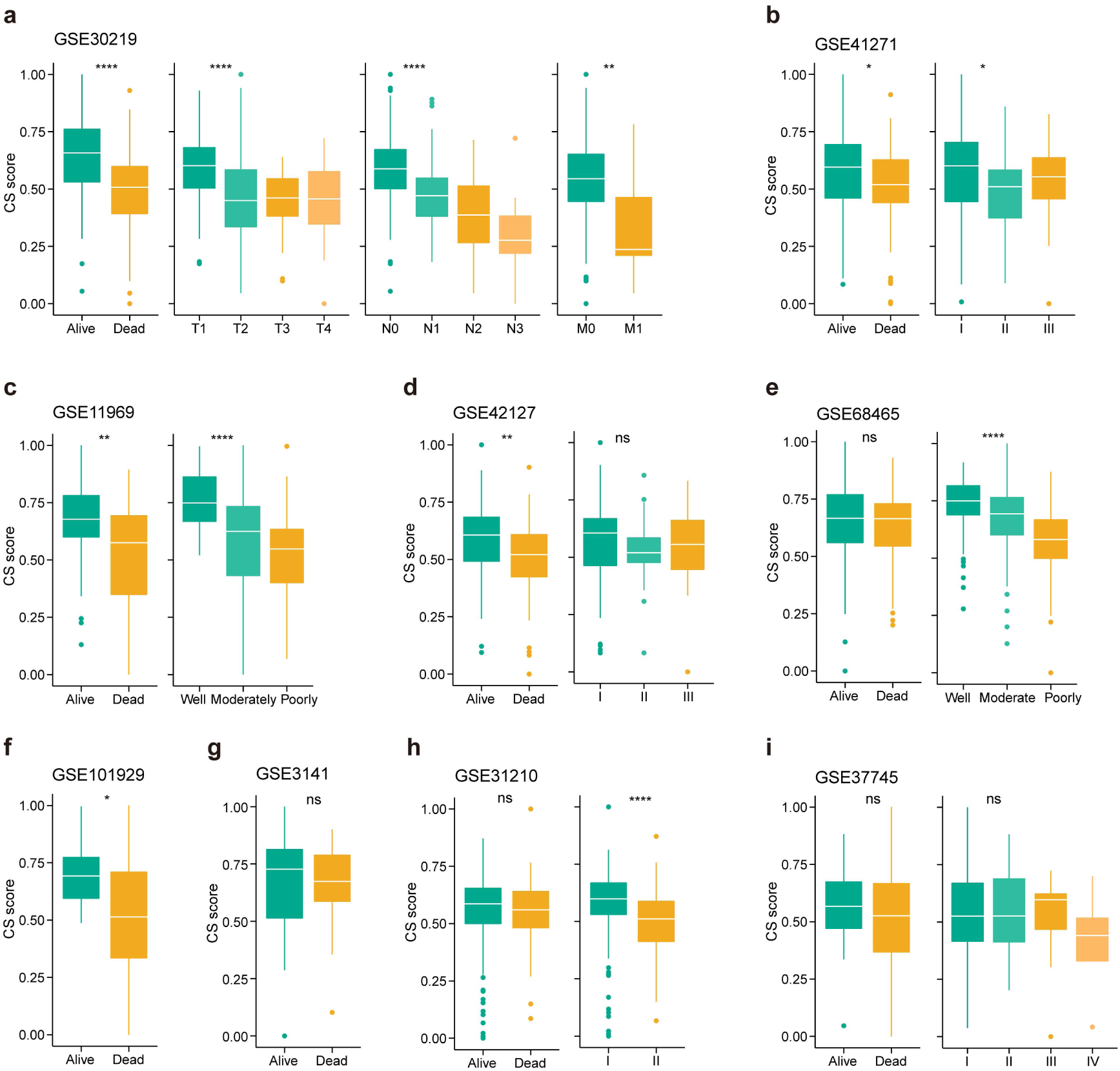
Cellular senescence (CS) predicts clinical indices across multiple lung adenocarcinoma (LUAD) datasets. **a–i** Box plots displaying the CS score in relation to vital status, American Joint Committee on Cancer (AJCC) and TNM stages in LUAD patients from the Gene Expression Omnibus databases, including GSE30219 (a), GSE41271 (b), GSE11969 (c), GSE42127 (d), GSE68465 (e), GSE101929 (f), GSE3141 (g), GSE31210 (h), and GSE37745 (i). Wilcoxon test *P*-values are presented to compare the two groups. Kruskal–Wallis test *P* values are presented for comparisons between more than two groups. \*\*\*\**P* < 0.0001, \*\*\**P* < 0.001, \*\**P* < 0.01, \**P* < 0.05, and ns: *P* > 0.05.

**Extended Data Fig. 4.**
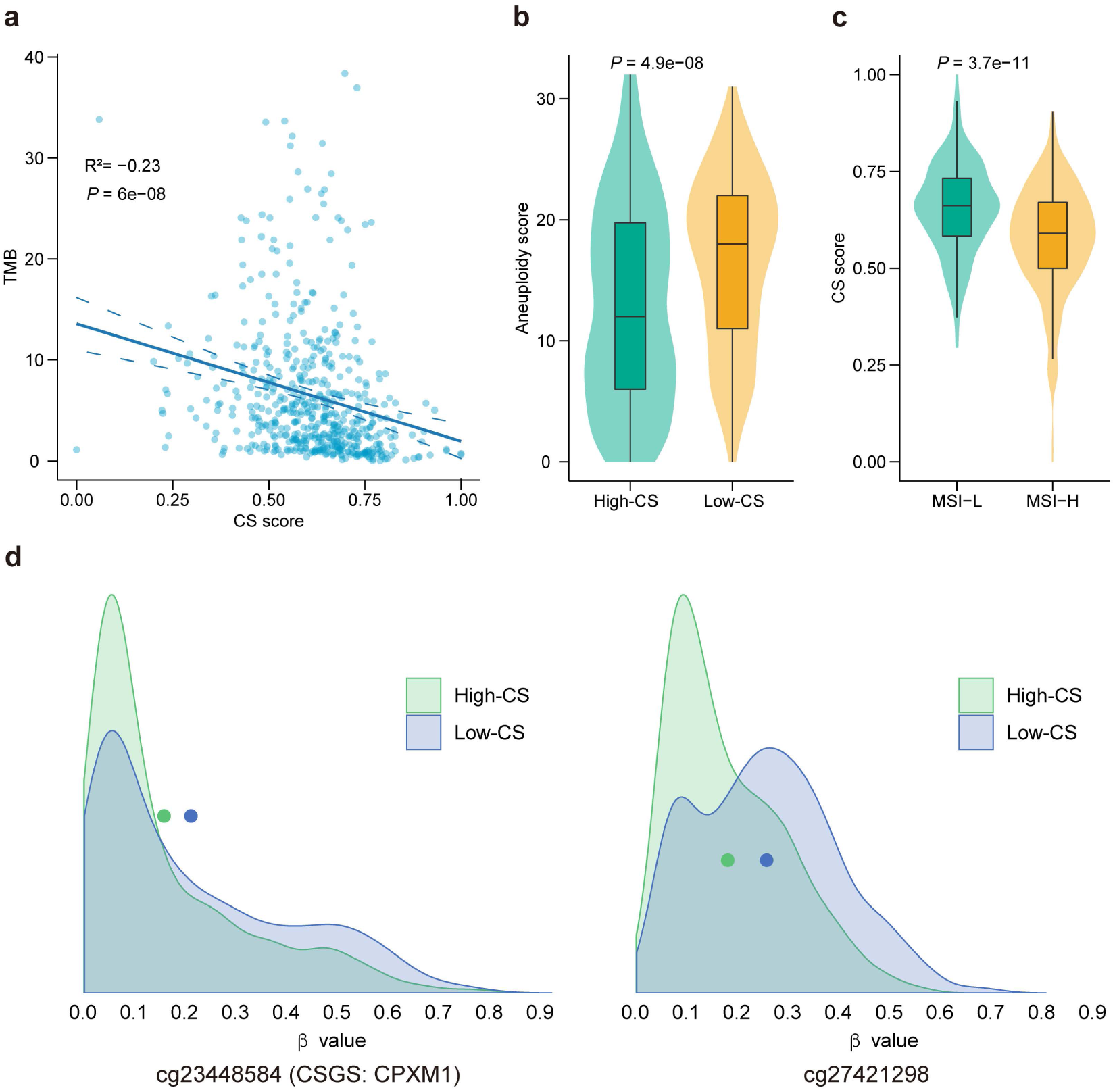
Cellular senescence (CS) related to multiple molecular characteristics in The Cancer Genome Atlas (TCGA)-lung adenocarcinoma (LUAD) dataset. **a** Scatter plots showing the correlation between tumor mutation burden (TMB) and CS scores in TCGA-LUAD. Statistical significance and correlation coefficients were determined using a rank-based Spearman’s correlation. **b** Violin plot showing the distribution of aneuploidy scores between the two CS subgroups. **c** Violin plots showing the distribution of CS scores between the two microsatellite instability (MSI) subgroups. Wilcoxon test *P* values are presented. **d** Ridge plot showing the distribution of estimated β values of two significant probes between the two CS subgroups based on the LUAD database, including cg23448584 (CPXM1, left) and cg27421298 (right). Dots in the ridge plot represent the median values for each group.

**Extended Data Fig. 5.**
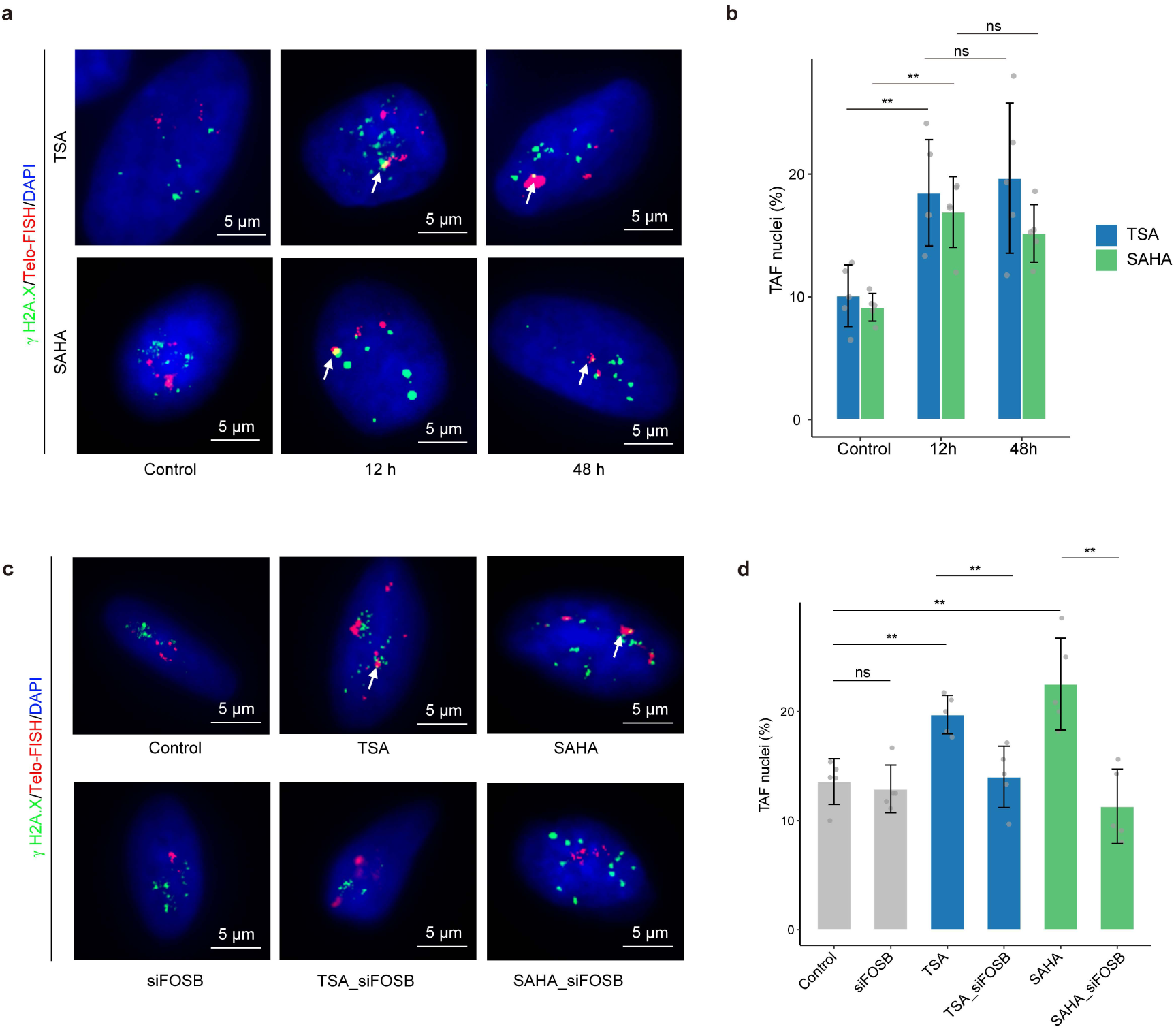
Validation of cellular senescence via Telomere-Associated Foci (TAF). **a, b** Representative images (a) of TAF staining in A549 cells (γH2A.X, green; Telo-FISH, red; DAPI, blue) treated with TSA (300 nM) or SAHA (500 nM) at different time points, and quantification of stained cells (b); **c, d** Representative images (c) of TAF staining in A549 cells (γH2A.X, green; Telo-FISH, red; DAPI, blue) subjected to different treatments, including control, siFOSB, TSA (300 nM), SAHA (500 nM), TSA with siFOSB, and SAHA with siFOSB, for 12 h, and quantification of stained cells (p). *n* = 5 biologically independent samples. Scale bar, 5 μm (Student’s t-test), and only the common groups with significant differences are shown. Data are represented as the mean ± SD. \*\**P* < 0.01, \**P* < 0.05, ns *P* ≥ 0.05.

## Supplementary table legends

Supplementary Table 1: Normal senescent cell lines

Supplementary Table 2: Tumor senescent cell lines or tissues

Supplementary Table 3: Publicly available cellular senescence signatures

Supplementary Table 4: Ranking outcomes for drug prediction

Supplementary Table 5: FOSB small interfering RNA (siRNA) sequences

